# Systematic review of biodiversity monitoring shows gradual rise of scaleable and automated methods and inherent spatial and taxonomic biases

**DOI:** 10.1101/2025.10.17.682377

**Authors:** Jenna Lawson, Cecilia Lee-Grant, Andrew O’Neill, Edward Forey, Lucas de Ruig, Thomas Weeks, Jingyi Yang, Robert Ewers, James Rosindell, William D. Pearse

## Abstract

Global policy efforts to reverse biodiversity declines are hindered by a lack of clarity over how to measure biodiversity. The debate over the best ways to define biodiversity and which metrics to use has taken place in the absence of a clear, holistic view of how biodiversity has been, and is currently, measured. This gap has hindered us moving from discussion to action on conserving biodiversity. Here we track trends in biodiversity measurement, using a dataset of *>*2400 papers across the last 22 years, identifying trends, biases, and gaps in the assessment of nine major taxonomic groups (Invertebrate, fish, aquatic mammal, terrestrial mammal, bird, herpetofauna, fungi, microbe and plant). We found proportional declines in invertebrate, but increases in fish, surveys over time, and fewer assessments of biodiversity in Africa and Oceania. Although we do find an increase in the use of high-throughput, scaleable methods such as eDNA and camera-traps, their uptake is more modest than might be expected and shows signs of having already plateaued. We argue that emerging technologies can, but remain underutilised in addressing the systematic biases in our understanding of biodiversity, and so in helping guide and track our progress towards restoring and saving biodiversity.

## 2 Introduction

Biodiversity is declining globally, with various sources suggesting a 69% average population decline (Almond, R.E.A. et al. 2022) and over a million species at risk of extinction (IPBES 2019). The rates of biodiversity loss likely vary across space and taxonomic groups (Vellend et al. 2013; Newbold et al. 2015; Leung et al. 2020), and biases in how data are collected across these axes impact the evidence-base of biodiversity-loss (Leung et al. 2020). Due to these biases, we still lack the evidence to understand the trends of both some of the most charismatic species (Junker et al. 2020) and our most threatened ecosystems (Maasri et al. 2022). Without this clear evidence base, calls to preserve biodiversity are beginning to be operationally framed around targets that aren’t biodiversity, such as abiotic proxies related to habitat amount and condition (Sayer et al. 2025). The agreement of the Kunming-Montreal Global Biodiversity Framework (GBF) by the Parties of the Convention on Biological Diversity (CBD 2022) and the establishment of the Intergovernmental Platform on Biodiversity and Ecosystem Services (IPBES) (Larigauderie and Mooney 2010) have been important steps towards assessing the state of biodiversity and international conservation agreements. However, their success relies on standardised monitoring methods that produce the data and evidence to quantify biodiversity change at the global scale and identify drivers of change (Navarro et al. 2017; Gill, M et al. 2017). How we effectively measure biodiversity to provide the data and subsequent evidence that feed into these frameworks is something that has yet to be defined or standardised.

We have made great progress towards defining biodiversity metrics, but that progress hasn’t been matched by clarity on how to collect the data to build the metrics. The CBD biodiversity indicators were designed to summarise biodiversity in a way that is useful for policy-makers (CBD 2010), but do not define how data should be collected or measured (Pereira et al. 2013). The Essential Biodiversity Variables (EBVs) defined under the Group on Earth Observations Biodiversity Observation Network (GEO BON) (Scholes et al. 2012), were designed to provide a general framework for biodiversity monitoring to support decision-makers (Navarro et al. 2017). They were developed specifically to facilitate data collection that uses standard formats and methods to ensure usability across datasets and therefore efficient use of evidence to guide conservation strategy and inform policy (Navarro et al. 2017; Pereira et al. 2013). There is hope that data collected for EBVs could then feed into indicators such as the Living Planet, Wild Bird, and Red List indices that are then used in policy frameworks (Navarro et al. 2017).

GEO BON has made major progress in defining EBVs and developing the framework around EBVs related to satellite remote sensing (Navarro et al. 2017). This is due to the availability of standardised data collection methods, such as Landsat images created by the Global Forest Change project (Pistón, N et al. 2023), which can then be used to feed into EBVs on ecosystem extent and fragmentation. EBVs that rely on in-situ observations (*i.e*., locally collected and not remotely-sensed) have made less progress, due partly to the challenges faced from lack of consistency around data collection methods and observations (Navarro et al. 2017). Understanding how we have and are measuring biodiversity across different taxanomic groups will aid us in developing standard practices going forward and providing consistent and comparable data to feed into metrics and measures developed to fill the evidence base.

There exist databases (GBIF 2024) and mobile apps (iNaturalist 2024) to collate observations of species, which can then be used to quantify biodiversity. Hundreds of biodiversity metrics have been proposed to quantify biodiversity (Purvis and Hector 2000; Santini et al. 2017; Marshall et al. 2020) and there is an ever-increasing desire to find a single metric to quantify biodiversity (European Commission 2014), yet how we effectively measure biodiversity to provide the data and subsequent evidence that feed into these metrics is yet to be defined. There also exists a wealth of literature detailing how to conduct surveys (Boyd, Bowen, and Iverson 2010; Hoffmann et al. 2010) and examining the efficiency of these methods (Doan 2003; Whitworth et al. 2017; Enari et al. 2019; Crunchant et al. 2020), yet an understanding of how and where methods are being applied and how this is changing overtime is largely non-existent. This lack of knowledge and consistency around data collection methods has held back the development of EBVs related to in-situ monitoring (Navarro et al. 2017)

With the landscape on biodiversity surveying and measurement changing and expanding rapidly, it is more important than ever to understand and standardise how we measure biodiversity. The availability of new technologies has exploded in recent years, with advancements in areas such as digital cameras traps, passive acoustic monitors, satellite data, and autonomous aerial vehicles (UAVs) (Pimm et al. 2015; Marvin et al. 2016). UAVs can map areas at wide spatial scales and at sub-centimeter resolution, multi-spectral Landsat imagery can measure biodiversity through vegetation and water, and networked devices can communicate, transfer, and process data across landscapes (Allan et al. 2018). The availability of new technology is truly impressive, and ten years ago it was suggested that developments in biodiversity monitoring technology would improve uptake within field-based survey methods (Pimm et al. 2015). Yet experts have not quantified how and where technology is being applied, nor have we standardised how they should be applied going forward in combination with observational approaches, as a way to quantify biodiversity to feed into metrics such as those required by the nature credit industries or at a policy level. If used correctly and embraced, these technologies have the ability to help address key conservation challenges (Pimm et al. 2015; Allan et al. 2018).

Here, we survey *>* 2400 papers since 2000 to establish an evidence base of how biodiversity has been measured and defined. This assessment describes historic and current gaps in knowledge and specific biases, how a range of methodologies are being used to measure biodiversity, and whether there has been an uptake of technology in the field as expected. This baseline is necessary to establish practical guidelines for biodiversity monitoring and brings us a significant step closer to closing the information gaps that hinder the establishment of key indicators and metrics of biodiversity.

## 3 Results

### Frequency change and popularity

We collected data on fourteen methods used to monitor biodiversity data across nine taxonomic groups (Fig. 1a and 1b) across 22 years (see Methods section for details).

**Figure 1.**
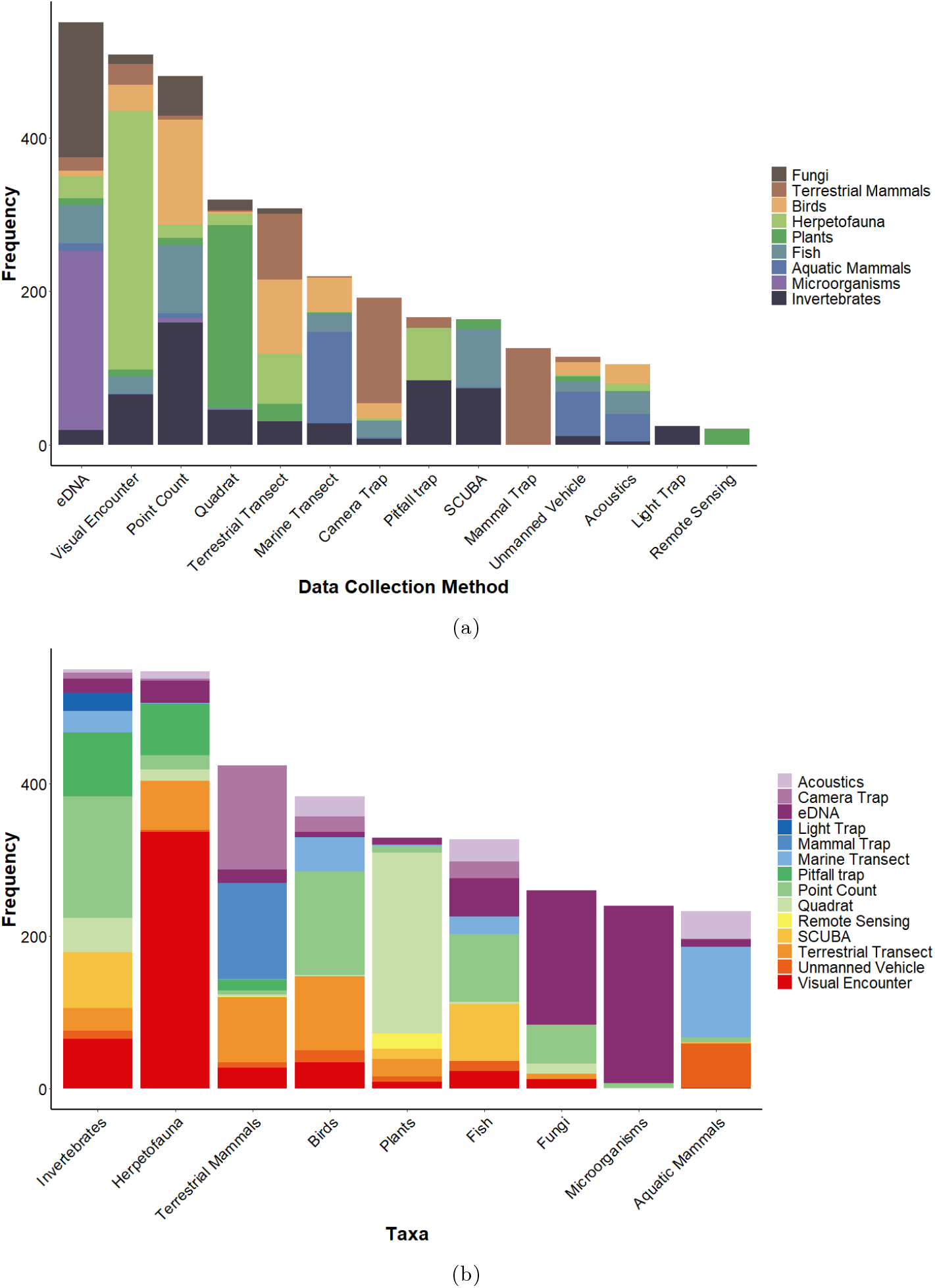
The diversity of taxa represented within each (a). Monitoring method and (b). Taxonomic group. The horizontal axis displays **(a).** the 14 methodological approaches used in this analysis and **(b)**. the nine taxonomic groups, presented in descending order of frequency (frequency is on the vertical axis). Each stacked section and of the bar represents an individual **(a)**, taxon or **(b)**. monitoring method, with representative colours detailed in the legend. **(a)**. Highlights both the commonality of each method for the study of biodiversity and whether its use is even across taxonomic groups. Some methods were found to have low diversity and evenness (*D* = 1), (*e.g*., light and mammal trapping), as seen by their dominance by one taxon and other methods exhibited higher richness and evenness (*e.g*., quadrats (*D* = 0.19), visual-encounter surveys (*D* = 0.21) and eDNA (*D* = 0.22)), as seen by their use across multiple taxon. **(b)** Highlights which methods we are using to study each taxa and how even this is. Some taxa were found to be very uneven (*D* = 0.33 −0.35), (*e.g*., microorganisms and fungi), as seen by their dominance by one method, whilst all others were even (*D* = 0.25 − 0.16), as seen by the use of multiple methods for their study.

We quantified the number of taxonomic groups that are represented within a method and how evenly taxonomic groups are represented within each method using simple richness measures (i.e. the count of papers using each method and taxa) and Simpson’s Evenness Index (*D*), where a score of 1 indicated low evenness of taxa or methods (Supp. Table 1). Perhaps unsurprisingly, we find that some methods are not diverse in use, only being used with very particular taxonomic groups (*e.g*., light trapping are used solely for invertebrates and mammal trapping solely for mammals), while other methods are popular across a wide range of taxa (*e.g*., quadrats, visual-encounter surveys and eDNA (environmental DNA); (Fig. 1a and Supp. Table 1). We also calculated the number of methods used to study each taxonomic group and how evenly spread methods are within a taxonomic group and found some taxa to be dominated by few methods (*e.g*., microorganisms and fungi are mostly collected using eDNA), while others were studied using a broad range of methods (Supp. Table 2 and Fig. 1b). eDNA methods have been in use for decades, and are the most popular kind of survey method (16% of all studies used eDNA). Despite showing high diversity, eDNA was dominated by just two taxa, with 32% of studies being related to fungi and 42% micro-organisms (Fig. 1a).

Poisson regression measured with a quadratic term for year showed support for a significant non-linear increase in publications through time (Supp. Table 3 and Fig. 2). We looked at the rate of change across all scientific publications, which has also increased through time, but not at the same rate. We find a ten-fold increase in the rate of annual biodiversity study publication in 2020 in comparison with 2000, even when background publication rate increases are accounted for (see Methods for details). General scientific studies increased by 188% and biodiversity studies from our study by 1818%. (taken from World Bank data; World Bank 2024). We do not compare past 2020 as world bank data end here.

**Figure 2.**
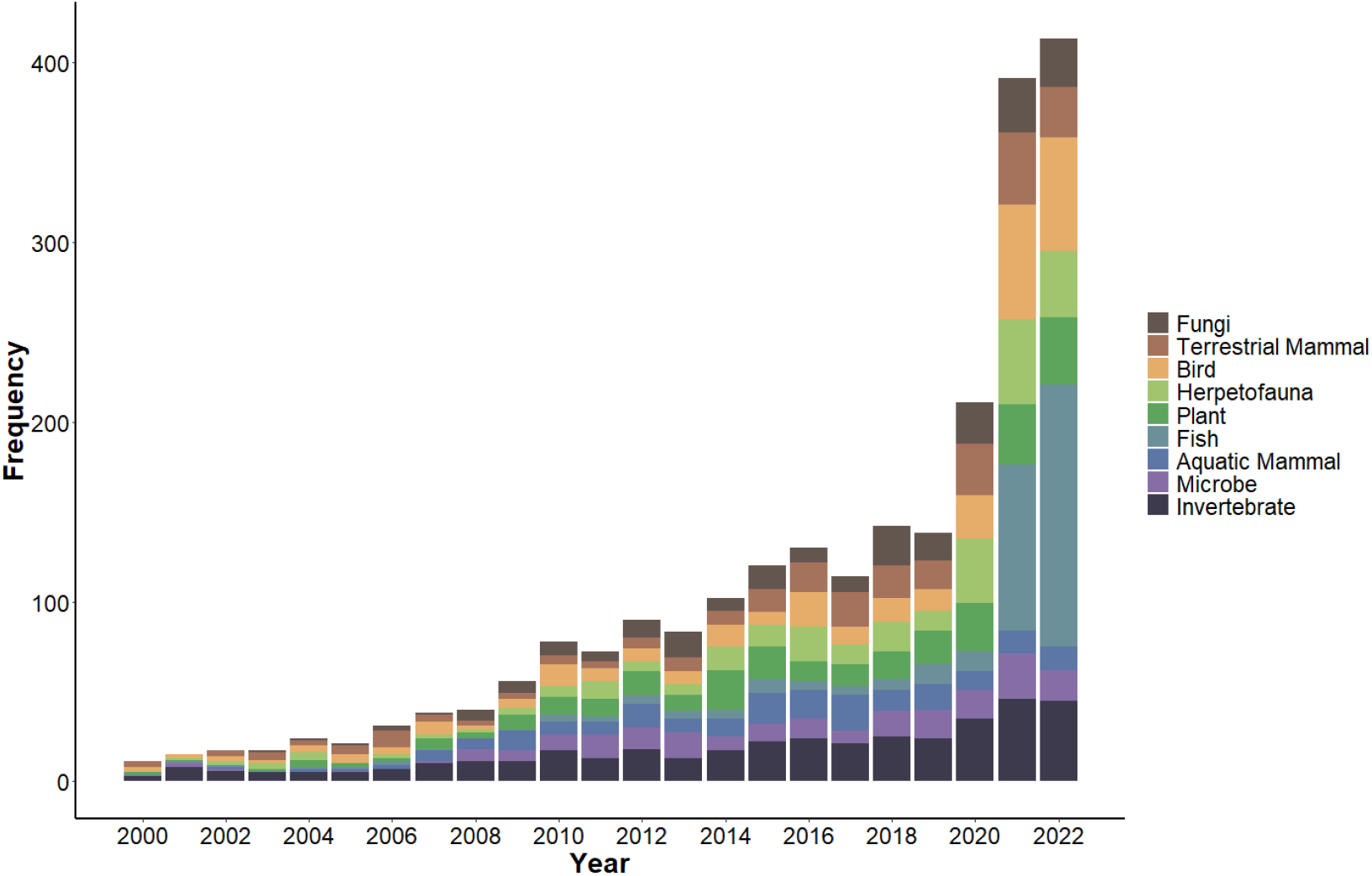
Frequency of published studies from 2000–2022. The horizontal axis displays the year 2000–2022, presented in ascending order of year (frequency of studies within a year is on the vertical axis). Each stacked section and of the bar represents an individual taxa with representative colours detailed in the legend. The figure highlights the rapid increase in the frequency of studies from 2020. Poisson regression with a quadratic term for year showed a significant nonlinear association between the publication of biodiversity studies through time (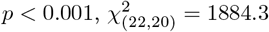, pseudo-*r*^2^ = 0.88).

### Taxonomic biases

While, in absolute terms, the number of biodiversity studies being conducted is increasing (Fig. 2), the proportion of that effort is not being spent equally across taxonomic groups. Binomial regression showed that studies of invertebrates are significantly declining through time (Fig. 3A and Supp. Table 4). Regression with a quadratic term for year showed a significant curved relationship between the proportion of studies on fish and time, with a rapid increase in studies after 2020 (Fig. 3B and Supp. Table 7). We also found a significant curved relationship between time and proportion of studies on birds (*p <* 0.001, *χ*2_(22,20)_ = 17.6, pseudo-*r*^2^ = 0.88) (Supp. Fig. 1A and Supp. Table 5) and aquatic mammals (*p <* 0.001, *χ*2_(22,20)_ = 52.91, pseudo-*r*^2^ = 0.88) (Supp. Fig. 1B and Supp. Table 6). The proportion of studies on birds declined until 2020, where we see a small increase, where as aquatic mammal studies increased until 2010 and have since started to decline. All other taxa are increasing or decreasing at rates that are essentially indistinguishable due to varying measures annually, with no significant difference when compared to the null model. Although not significant we do note a decline after 2006 in the study of a traditionally oversampled group, terrestrial mammals (*p* = 0.24, *χ*2_(22,21)_ = 1.35, pseudo-*r*^2^ = 0.21) (Supp. Table 17). Notably, this means that our tracking of the most diverse taxonomic group on Earth (invertebrates) is declining through time, suggesting the systematic under-representation of these groups is increasing.

**Figure 3.**
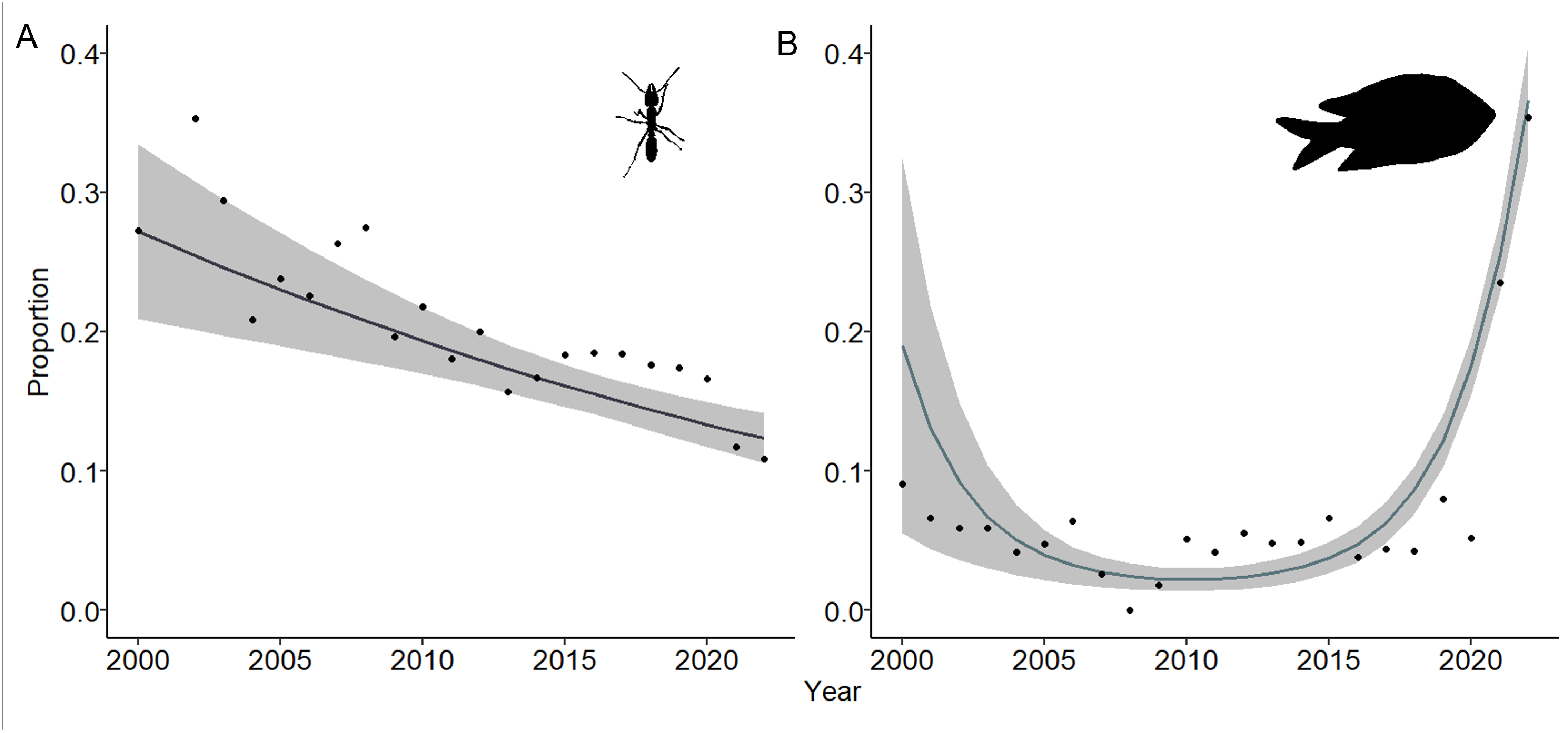
Proportional change in published studies for key taxonomic groups overtime. The horizontal axis displays years from 2000–2022. The vertical axis shows the proportional change in the number of studies for each taxonomic group, with year on the horizontal axis. Of the nine taxonomic groups included in the meta-analysis we display here only the key results for invertebrates and fish. Binomial regression showed a significant decrease in the proportion of studies on invertebrates (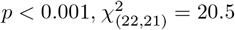, pseudo-*r*^2^ = 0.75) overtime and B. Polynomial regression shows a significant curved relationship between time and proportion of studies on fish (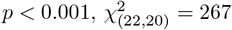, pseudo- *r*^2^ = 0.96), with a rapid increase from 2020.

### Changes in methods

We combined methods into high- and low-throughput to look at change in throughputness over time. Using binomial regression we see significantly higher use of low throughput methods across studies between 2000 and 2022 (*p <* 0.001, *χ*2_(45,44)_ = 207.52, pseudo-*r*^2^ = 0.83) (Supp. Table 28). Yet over time, binomial regression with a quadratic term for year shows a significant increase in high-throughput studies compared to low-throughput (*p <* 0.001, *χ*2_(22,20)_ = 26.419, pseudo-*r*^2^ = 0.88) (Fig. 4D and Supp. Table 27). Despite an explosion in the availability of high-throughput technologies for the study of biodiversity (see methods section for classification of high- and low-throughput methods), only the use of camera-traps and eDNA have seen a significant change over time. Binomial regression showed that the proportion of camera trap studies have gradually increased overtime (Fig. 4A and Supp. Table 14), while binomial regression with quadratic term for year highlighted a curved relationship between the proportion of eDNA studies and time, increasing until 2013, where studies begin to level off (Fig. 4B and Supp. Table 15). Autonomous-vehicles and remote-sensing showed no statistically significant trends through time, but we only detected their use since 2004 and 2009, respectively (see Supp. Material Fig. 2). The proportional use of low-throughput in-person, observational survey methods varies between methods. The proportion of studies using pitfall trapping (*p <* 0.001, *χ*2_(22,21)_ = 13.488, pseudo-*r*^2^ = 0.83) and quadrats (*p <* 0.001, *χ*2_(22,21)_ = 5.6556, pseudo- *r*^2^ = 0.53) have experienced a gradual decline in use (Supp. Table 19 and 21), while the proportion of studies using point counts showed a significant curved relationship, gradually declining until around 2020, where studies begin to increase, however further data may be necessary to confirm this upward trend (*p <* 0.05, *χ*2 = 25.35_(22,20)_, pseudo-*r*^2^ = 0.79) (Supp. Table 20). We also found a significant curvilinear relationship between time and proportion of studies using marine transects (*p <* 0.001, *χ*2_(22,20)_ = 26.589, pseudo *r*^2^ = 0.80), with an increase until 2010, followed by a gradual decline (Fig. 4 and (Supp. Table 18). While this could be in part related to the proportional decrease in marine mammal studies, other common methods that are decreasing, such as terrestrial transects and visual encounter surveys, are common across a range of taxa, many of which are increasing or showing little change overtime. The exception to this is SCUBA studies, which have shown a significant increase in use (Fig. 4 and Supp. Table 23).

**Figure 4.**
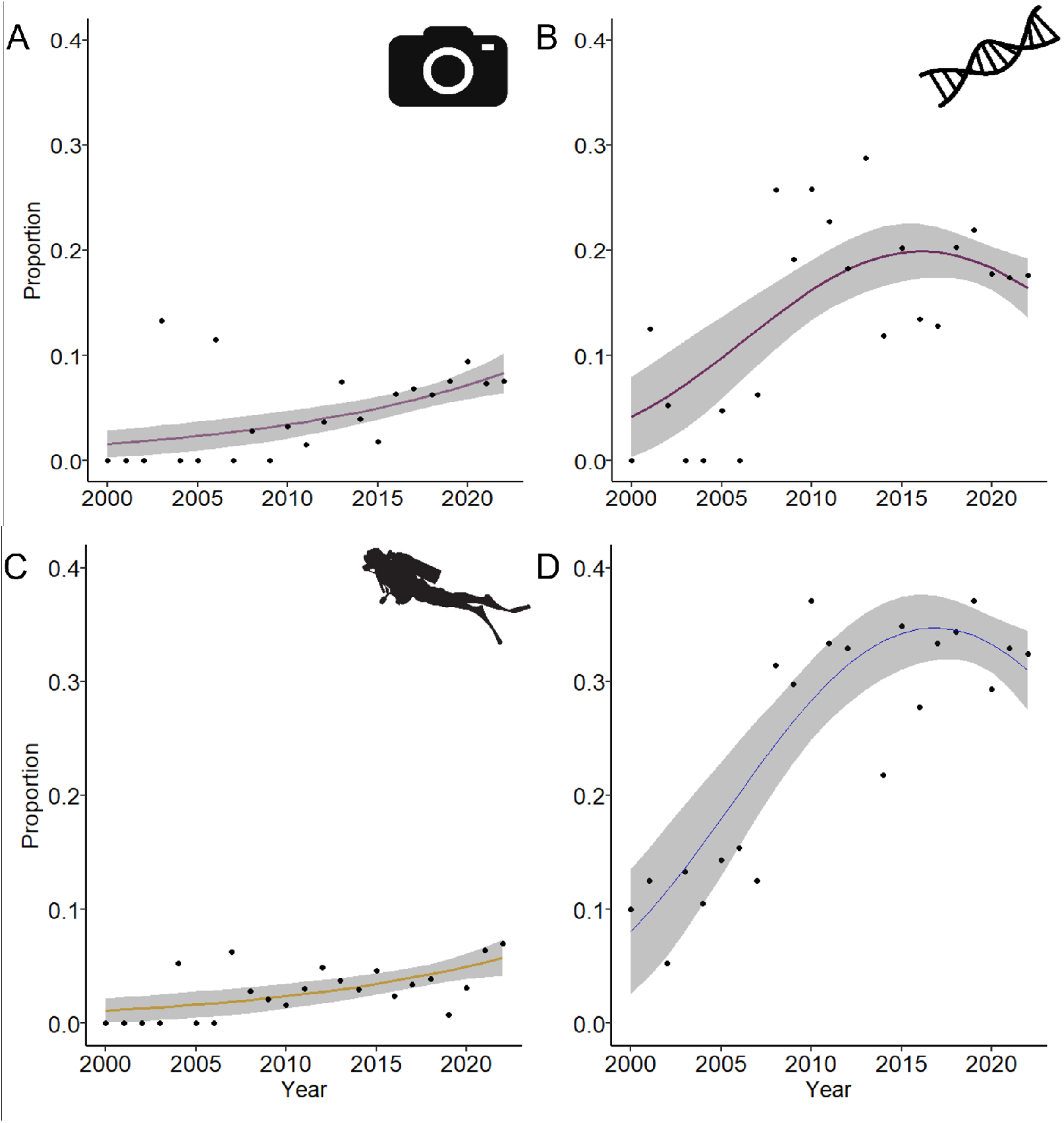
Proportional change in published studies for key monitoring methods and throughputness overtime. The vertical axis displays the proportional change in the number of studies for each each method and throughputness (year, from 2000–2022 is on the horizontal). Of the 14 methods included in the meta-analysis we display here only the key results for methods increasing in use: camera traps, eDNA and SCUBA and high throughputness. Binomial regression showed a significant increase in the proportion of studies using A. Camera traps (*p <* 0.001, *χ*2_(22,21)_ = 15.27, pseudo-*r*^2^ = 0.88) and C. SCUBA overtime (*p <* 0.01, *χ*2_(22,21)_ = − 10.386,pseudo-*r*^2^ = 0.88). Binomial regression showed a significant curved relationship between time and proportion of studies using eDNA (*p <* 0.001, *χ*2_(22,20)_ = 14.315, pseudo-*r*^2^ = 0.88), with a rapid increase from 2000, leveling off around 2020. D. We see a significant increase in high-throughput studies until 2010, where this change then levels off (*p <* 0.001, *χ*2_(22,20)_ = 26.419, pseudo-*r*^2^ = 0.88).

### Geographic and Biome Biases

Chi-square goodness of fit test confirmed that studies are not evenly distributed across biomes. There was a difference between percentage of studies across all biomes, with 16% more studies in the marine biome compared to freshwater, 36% higher in the terrestrial biome compared to freshwater ecosystems and 75% higher in the terrestrial compared to the marine. In total 58% of studies took place in the terrestrial biome, 26% in marine and only 16% in freshwater ecosystems (Fig. 5). Chi-square goodness of fit test also confirms that studies are not evenly distributed across continents. With the exception of Antarctica, Africa and Oceania were studied the least with just 8% and 6% of all studies respectively. South America sits somewhere in the middle with 15% of all studies, with Europe, North America and Asia holding the most, with 18% 22% and 29% respectively (Figure 6). Whilst these results may be somewhat due to the size of continents this certainly isn’t the case for Africa, as the 2nd largest continent after Asia.

**Figure 5.**
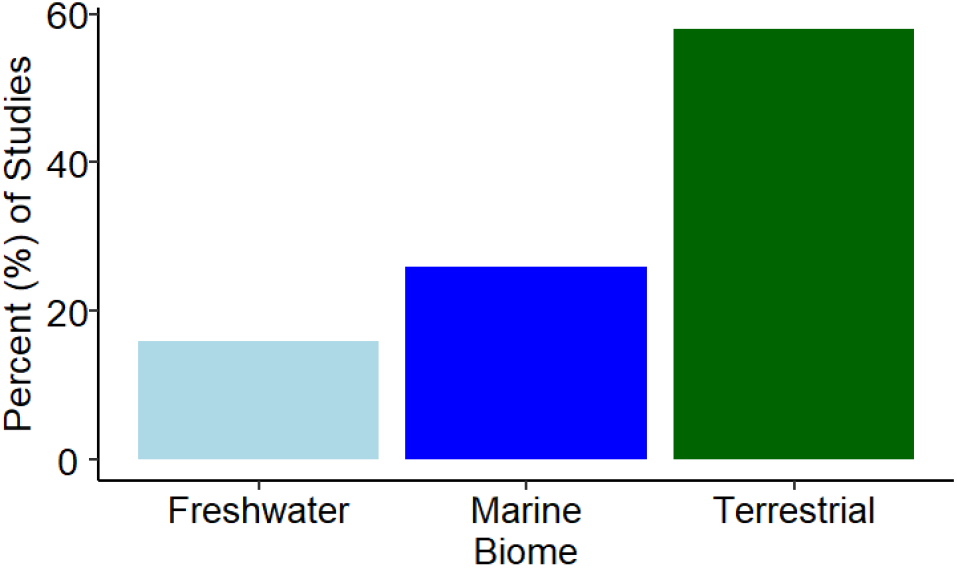
Frequency of studies per biome. The horizontal axis displays the three biomes, with categories sorted in ascending order order by percent of studies. The vertical axis shows the percent of studies for per biome. Chi-square goodness of fit test shows that studies are not evenly distributed across biome (*p <* 0.001, *χ*2_(2)_ = 691.28). There were more studies in the marine and terrestrial biomes compared to freshwater and in the terrestrial compared to marine.

**Figure 6.**
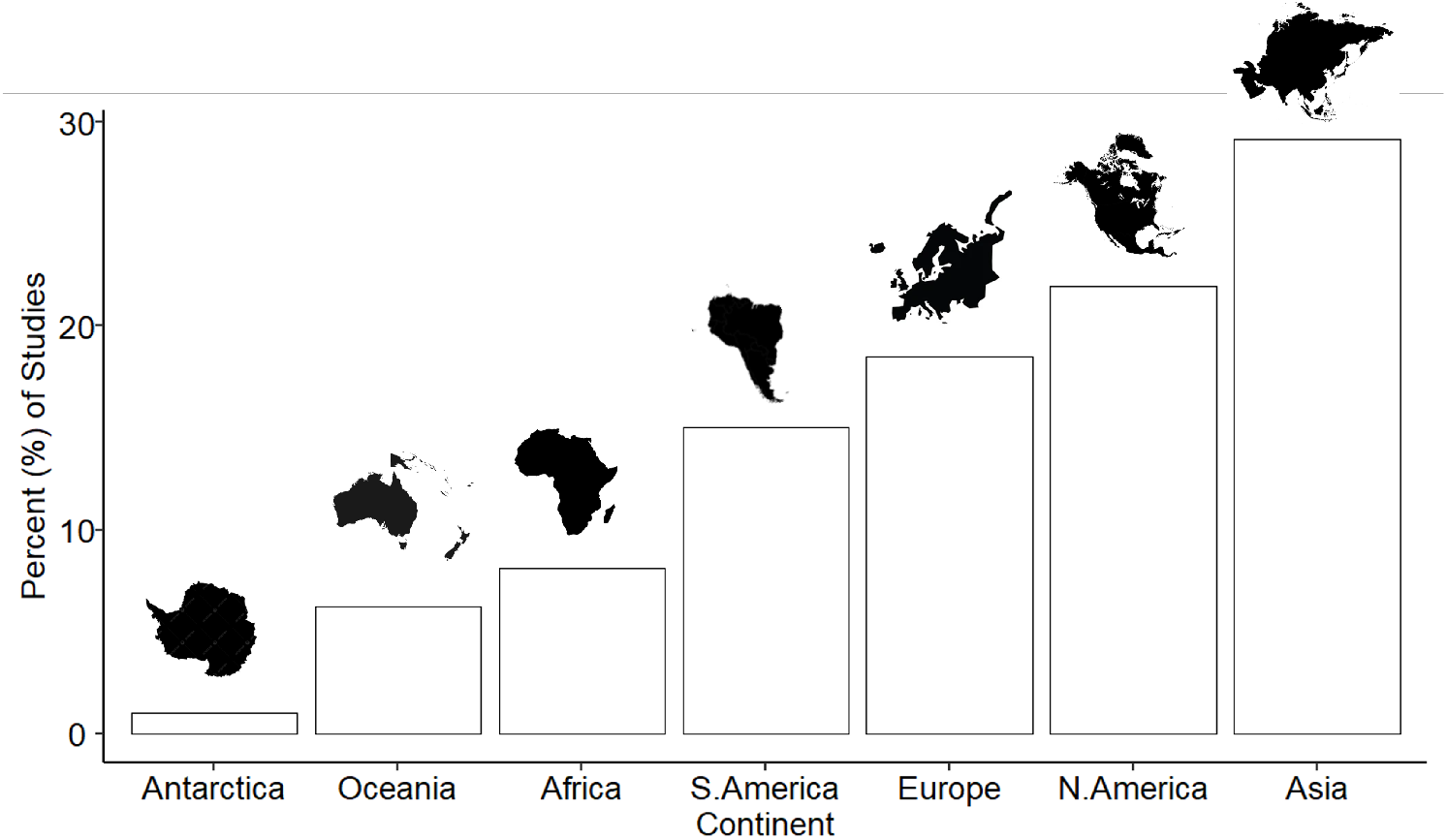
The frequency of studies for each continent. The horizontal axis displays the seven continents, with categories sorted in ascending order order by percent of studies. The vertical axis shows the percent of studies for per continent. Chi-square goodness of fit test confirms that studies are not evenly distributed across continents (*p <* 0.001, *χ*2_(2)_ = 958.15). North America, Europe and Asia hold the most studies, whilst Antarctica, Oceania and Africa hold fewer studies.

## 4 Discussion

This meta analysis surveyed *>* 2400 papers over 22 years to assess how a range of methodologies have been used to measure biodiversity and define gaps in knowledge and specific biases. Whilst the number of studies of biodiversity has increased, the proportion of effort is not being equally spread across taxa, with significant decreases in the proportion of invertebrate studies. We also find a significant under-representation of studies in the freshwater biome and a strong geographic bias, with Africa and Oceania receiving little attention across the literature. We see a modest increase in the use of high-throughput methods, but technologies that have been available for over a decade are likely not being used to their full potential.

### 4.1 The rapid increase in studies

The increase in published literature over time highlights the increasing popularity of biodiversity, likely related to an increased urgency to quantify and mitigate anthropogenic impacts on biodiversity, and the increasing interest in citizen science (Chowdhury et al. 2023). The rapid increase in published literature since 2020 could also be related to the onset of COVID-19, where data collection activities and new projects were put on hold for many researchers, allowing time to consolidate previous research and publish papers. The observed increase in biodiversity studies through time could potentially result from bias in our data collection method. We searched the literature for keywords, returned results based on relevance (which is likely influenced by publication year) and did not stratify by year. Nevertheless, we suggest our finding of an increase in publication rates is not spurious for three main reasons. First, the rate of change in biodiversity studies is not monotonic: the year on year percent change in publications declines in 2004, 2010, 2012, 2016, and 2018, suggesting it is not a constant sampling artifact. Second, publications across all fields have seen an increase, just not at the same pace (World Bank 2024). Third, our findings are consistent with previous literature analyses that were limited to camera traps (Delisle et al. 2021) and eDNA studies (Beng and Corlett 2020).

### 4.2 Informing standardisation

Our meta-analysis highlights the variability in methods used to monitor different taxa. Some methods, such as light- and mammal-traps, are predominantly used for and suited to one or two taxa, whilst others, such as visual encounter surveys and quadrats are popular across a range of taxa. Microorganisms are almost solely studied using eDNA, potentially indicating that this is the best monitoring method for this group. Practically, this makes sense as microorganisms are unlikely to be studied via other approaches that are less well suited to them. Aquatic mammals are predominately monitored using SCUBA, acoustic and marine transect surveys, therefore a combination of these methods may be the best approach for studying this group. However some taxa, such as fish and birds, use a range of monitoring methods, showing a clear lack of any standardised approach. Lack of standardisation within study methods can limit applicability and comparability in certain circumstances (**napier·advancements·2024**), such as the need to quantify change in biodiversity to feed into policy or nature credit metrics. In attempting to define standardised approaches to monitoring, the evidence here shows that for some taxa, only one method may be required for effective study, however for others, a suite of methods may be required, making standardisation more challenging. By looking at how we have been studying different taxa and the range of methods we have used, this provides an excellent starting point for considering what the most appropriate methods might be going forward. Using evidence of what approaches have been used and considering the current advances in the field, we suggest that taxonomic experts attempt to define a set of methods that should be used for each taxonomic group, ensuring outputs can feed into metrics used within conservation agreements in a standardised way.

### 4.3 The rise of technology

Technology is changing the way we study biodiversity, and whilst this is essential to improve the scale at which we can study, it is complicating the standardisation of monitoring approaches as there are ever more options available. Pimm et al. (2015) suggested that the uptake of technology for the study of biodiversity had been slower than expected, but expected uptake to increase. Our data suggests that higher-throughput methods (UAV’s, remote sensing, camera traps, eDNA and acoustics) all showed a proportional increase in use, but this trend is only significant for camera traps and eDNA. For camera traps this is likely related to the rapid development in hardware in the early 2000s and their application to terrestrial mammals (Oliver et al. 2023), a traditionally over-sampled group. With improvements in hardware and machine learning algorithms for analysis, we are increasingly seeing camera systems being used for the study of other taxonomic groups, especially birds (Oliver et al. 2023), invertebrates and fish (Roy et al. 2024; Mallet and Pelletier 2014).

The extraction of DNA from environmental samples can characterise communities from a range of taxa (Venter et al. 2004; Ficetola et al. 2008) and is thought of as an emerging and novel approach due to its ability to rapidly quantify biodiversity at scale (Ruppert, Kline, and Rahman 2019; Beng and Corlett 2020), yet we find that it has been used for the study of microbes and fungi since 2000 and likely long before this. This suggests that eDNA is only considered emerging in some fields, likely due to a historic absence of reference libraries and availability of alternative approaches (Ruppert, Kline, and Rahman 2019). In a study of literature between 2008-2018 eDNA was found to be increasing in popularity, however the authors did not consider microorganisms and fungi as separate categories (Beng and Corlett 2020). By doing so here it may explain why we find its use levelling off and not increasing exponentially. By not considering the use of eDNA in the study of microorganisms and fungi it has lead the field to consider eDNA an emerging method, which it is not and we may have missed out on an important opportunity to transfer learning between taxonomic groups. By separating microorganisms and fungi in this study we have also shown that eDNA could be considered a standardised approach to the study of microoprganisms and fungi. As reference libraries continue to improve it is likely we will see increased use of eDNA within across taxonomic groups and mediums, such as the more recent airbourne eDNA for vertebrates (Lynggaard et al. 2022).

Despite an initial upward trend in the use of higher-throughput methods, we see a levelling off after 2010, suggesting that whilst there was an initial uptake during the first decade of the 21st century, that trend hasn’t continued exponentially in proportional terms. Lower-throughput approaches also remain more common at every year. There are likely numerous reasons for this, including unsustainable financing and inadequate capacity building (Speaker et al. 2022; McGeady et al. 2023; Reynolds et al. 2025), the requirement for benchmarking against traditionally used approaches (Borja et al. 2024), robustness of hardware, legal restrictions and impacts on animal welfare (MacWilliams, Kim, and Trueman 2024) and lack of expert labelled data for training machine-learning algorithms that are usually required in the analysis of big data (Rasmussen, Stowell, and Briefer 2024; Wägele et al. 2022). Barriers to uptake of technology were found to be more pronounced in low and middle-income countries (LMICs), due to constraints on development funding and upfront costs and for women, again due to constraints on development funding and their perceived technical skills (Speaker et al. 2022; McGeady et al. 2023). It is also the case that technology is predominately development in higher income countries, which is particularly concerning as much of the world biodiversity exists in LMICs (Reynolds et al. 2025). Integrating novel technologies into standardised monitoring frameworks will be a challenge until they can be more inclusive in their taxonomic coverage (Stephenson 2020) and until they are reliable, affordable, and capacity for use is widespread, especially in biodiversity-rich countries. Until then it is likely that any standardised approach to monitoring will require higher-throughput methods to be complemented with traditional observational approaches for the foreseeable future.

Despite the continued popularity of lower-throughput methods, observational approaches such as point counts, quadrats, marine transects and pitfall traps have declined, likely replaced or complemented in some part by higher-throughput methods. The decline in use of marine transects could be in part due to a similar drop in the study of marine mammals, however, advances in the field of aerial, surface and underwater vehicles, although lacking in applications to biodiversity, are showing significant promise for the study of marine mammals (Gooday et al. 2018; Kelaher et al. 2019). Reasons for a decline in pitfall trap studies could be due to traps indiscriminately killing many non target animals, especially small vertebrates (Zhao et al. 2013), and species and attractant biases which lead to unreliable estimations of diversity and density (Pearce et al. 2005). Whilst some observational approaches have declined in use since 2000, for good reason, they are clearly still widely used methods and have a place in the monitoring of biodiversity across taxa. As higher-throughput methods improve and hopefully become more accessible globally, this pattern may change. Of importance to note is that many methods are increasing through the throughputness scale as technological advances continue. For example, camera traps data that were once analysed manually, limiting scalability, is now largely automated through the use of machine learning approaches, which will likely accelerate its use globally (Vélez et al. 2023).

An inconsistency in the decline in observational studies is SCUBA studies, which have seen a gradual increase in use. We do not believe this is related to the increase in fish studies due to the difference in timing, with SCUBA seeing a gradual increase overtime and fish showing a rapid increase since 2020. Explanations for this increase could be due to the urgent need to monitor shallow water reefs as they become increasingly affected by bleaching events, which are now around five-times more common than they were 40 years ago (Hughes et al. 2018). It is also possible that the growing SCUBA diving industry is using their skilled workforce and citizen science to support marine studies and protect the ecosystems that their economy depends on (Goffredo, Pensa, et al. 2010; Goffredo, Piccinetti, and Zaccanti 2004; Hermoso, Narváez, and Thiel 2021; Nisa, Schofield, and Neat 2022).

### 4.4 Taxonomic and geographic bias

The disparity in our measurement of biodiversity is leaving some of the most diverse taxa and areas of the Earth understudied and a lack of evidence to feed into metrics. Invertebrates have been historically under studied (Clark and May 2002; Di Marco et al. 2017) and it is concerning to see that this trend is still continuing, despite calls to improve monitoring (Thomas, Jones, and Hartley 2019) and evidence of rapid declines globally (Lister and Garcia 2018; Hallmann et al. 2017). It is however encouraging to see that the proportion of fish studies are increasing overtime, as previous research found these taxa to be unrepresented within the literature (Clark and May 2002; Di Marco et al. 2017; Troudet et al. 2017). This could be in part related to the increase in SCUBA studies, as discussed above and the importance of the fisheries and aquaculture industries for the global economy (Meinam et al. 2023). We note that the increase in studies on fish is controlled by only two data points in 2021 and 2022, and whilst we do not believe these to be related to our study design, as we analysed a similar amount of fish publications compared to other taxa, this upward trend should be treated with caution. Despite the increased proportion of studies on fish, freshwater ecosystems remain under-represented in the literature, as has previously been found, (Trimble and Aarde 2012; Di Marco et al. 2017), despite being destroyed at twice the rate of marine or terrestrial ecosystems (Linke et al. 2018). Experts have called for more focus on freshwater ecosystems (Linke et al. 2018), something we still have not seen come through in the literature.

A historic under-representation has existed for studies across Africa (Trimble and Aarde 2012; Di Marco et al. 2017), a trend that we still see in our meta-analysis for Africa and indeed for Oceania. In a review of Africa’s progress towards the Aichi targets, it was found that huge areas of Africa remain undocumented for terrestrial mammals, birds and amphibians and the report suggests the same is likely to be the case for other lesser studied taxa (Farooq et al. 2021). It is suggested that this is due to researchers returning to previously mapped areas and not exploring new areas, lack of national resources and local capacity (Farooq et al. 2021). As discussed above, We will only correct our biases by improving accessibility to monitoring, building capacity among local scientists and improved funding. It should be noted that we excluded papers that were not in English, therefore this may have biased our results towards english-speaking countries.

Taxonomic, geographical and ecosystems scale gaps in scientific knowledge lead to inaccurate assess-ments of status, trends, threats and misleading conservation priorities (Di Marco et al. 2017; Farooq et al. 2021), making it essential to address these critical information gaps. If the aim is to develop standarsised monitoring frameworks it is essential that we are collecting the data and evidence required to feed into national and international metrics, policies and frameworks (Stephenson, Ntiamoa-Baidu, and Simaika 2020).

Its encouraging to see that studies monitoring biodiversity have rapidly increased in recent years, in line with the need to understand the scale of damage and develop mitigation strategies (IPBES 2019), but there is still a lot of work to be done. Until we can close critical data gaps geographically and across taxa and ecosystems, we will not be collecting the required data to feed into the variables, metrics and frameworks created to measure and monitor biodiversity. Future research should focus on understudied taxa (*e.g*., invertebrates), ecosystems (*e.g*., freshwater) and regions (*e.g*., Africa and Oceania) and in developing the technology that could monitor these areas at the scale required. This meta-analysis has shown how we have been measuring biodiversity, which can be helpful going forward in defining common, appropriate and standardised monitoring frameworks. The use of high-throughput methods such as eDNA and camera traps are increasing and diversifying in use across taxa and technology offers great promise to monitor biodiversity at the scale required to achieve targets such as those set in the Kunming-Montreal Global Biodiversity Framework. However, uptake has not increased as expected and continued innovation is required to tackle challenges related to hardware and analysis. For example, the data storage requirements represent a large financial and environmental cost, something which could be tackled through the continued development of edge computing, allowing only the key raw data to be saved. A wider challenge in ensuring uptake of technology centres around equitable development. For example, the majority of hardware and AI development is carried out in the global north, yet many of the earth’s most biodiverse regions are in the global south. Future work should focus around improving accessibility to and capacity to use technology at the national level, through local production of hardware, open science and capacity building, particularly for disadvantaged groups. The use of lower-throughput methods is still common and will likely remain so, especially until the challenges around higher-throughput methods are solved. Any standardised monitoring framework for biodiversity should include both approaches and should be seen as complementary and work should focus around the complementarity of these approaches.

## 5 Methods

### 5.1 Literature Search

The overarching aim of this analysis was to understand how field-based studies are measuring biodiversity and to analyse how this changes across spatio-temporal factors such as by time, geography and biome. Biodiversity encompasses all biological diversity on earth, from genes to ecosystems, and including all extant taxa. Therefore, capturing all of this information was complex. Methods used to study biodiversity are dependent on many factors, including financial and operational limitations, the country and habitat where the study is conducted, the specific aim of the study, the skill level and background of the personnel, and the taxa being studied. Since many of these factors are difficult to separate and quantify, we decided to investigate the measurement and quantification of biodiversity separately for different types of taxa: plants, terrestrial mammals, aquatic mammals, birds, invertebrates, fish, herpetofauna (assessing both amphibian and reptile taxa), fungi and microorganisms. These taxa were chosen to include the five major groups of vertebrates, splitting marine and terrestrial mammals due to the difference in monitoring approaches and the separate kingdoms of plants and fungi. We also chose to include microorganisms as they generally don’t fit into any of the categories above, due to their importance in the study of biodiversity and owing to their size we concluded that monitoring methods would be very different.

With so many methods used to measure and quantify biodiversity, it would have been challenging to capture detailed information on all methods. The design of the meta-analysis didn’t exclude any methods from being included. We chose 14 methods to carry through to the analysis stage, based on our knowledge and expertise of which methods were either common across taxa, or were emerging methods that we expect to become prominent in the near future. The methods chosen were points counts, terrestrial transects, marine transects, quadrats, pitfall and pan traps, light and malaise traps, small mammal traps, visual encounter surveys, eDNA, acoustics, camera traps, SCUBA surveys, remote sensing and unmanned vehicles. Where other methods were found in the literature review, these were recorded in a separate section to provide a full picture of methods used, however no further analysis of their use was carried out.

#### 5.1.1 Text-mining

To synthesise data for the meta-analysis while reducing the bias in key word selection, we performed a systematic search of the scientific literature using the R package `litsearchr` Grames2019. This package facilitates a quick, objective, reproducible search strategy, and uses text-mining and keyword co-occurrence networks to identify important terms to include in the final search. We performed one systematic search for each taxon.

The following process was used to synthesise papers for the meta-analysis for each taxa. Firstly, a precise Boolean search, known as the ‘naive’ search, was performed that returned a set of highly relevant articles. This was written for each taxon using the following structure: ‘taxon AND biodiversity AND survey’ (See Section **??**). The search terms were entered into Web of Science and Scopus for each taxon to generate a preliminary list of articles per taxa. For herpetofauna, most studies use the terms “amphibian”, “reptile”, and “anuran”. These were the specific terms used in the search, and were later combined into the higher level taxon group “herpetofauna” due to the incidence rate (38.3%) of studies assessing both groups simultaneously, and the similarities in biodiversity surveys across amphibian and reptiles studies.

`.RIS` files were then downloaded containing the key details for all articles in the search list. Information included in `.RIS` files included title, keywords and abstract. These `.RIS` files were imported into R through the `litsearchr` package and duplicated articles from Scopus and WoS were removed.

To extract potential keywords from articles in the de-duplicated dataset, the Rapid Automatic Keyword Extraction (RAKE) algorithm was used within `litsearchr`. This algorithm is designed to identify keywords in scientific literature by selecting strings of words uninterrupted by stopwords or punctuation. Within `litsearchr`, keywords were extracted if they occurred more than 100 times in the title or key words across all articles. Potential keywords were then used to generate a document-feature matrix using the potential keywords as features and the combined titles and keywords of each article as the documents.

From this, a keyword co-occurrence network was generated to measure each term’s importance and influence in relation to the topic being reviewed. In the network, each node represents a potential search term and the edges are co-occurrences of two terms in the title, abstract, or tagged keywords of a study. Node strength generally follows a power law, therefore, there are regions of rapid change where nodes become increasingly more important. To identify these regions and the points beyond which node strength increases dramatically, nodes were ranked by strength and a genetic algorithm was used to find rapid change points in keyword importance. To determine which keywords were to be considered for inclusion in the search terms, `litsearchr` takes the strength of the node at the first knot and retrieves the keywords associated with all nodes stronger than it. `litsearchr` then suggests the most appropriate and important words for the topic of study.

Suggested terms were then manually reviewed to determine which were appropriate for the final search string; terms that were too specific were not included. The final search strings were then entered into Web of Science and Scopus to generate a new list of suitable articles. This final search was further refined by article type to exclude fields that were not relevant, for example, parasitology or meteorology. Book chapters and reviews were not included to reduce the number of articles that did not collect field data. We reviewed all returned papers to determine their suitability for inclusion within the meta-analysis. Papers were excluded if they were not in English, if they did not use field-based methods to collect data and if methods were not described in detail. In total found 2444 suitable papers, broken down by taxa (Table 1). All papers returned by either WoS or Scopus, were in English and had conducted fieldwork to collect data on biodiversity. All suitable papers were included in the analysis.

**Table 1.** The number of papers included in the methods per taxa. Table showing the number of papers included in the methods per taxa

| Method | Number of studies |
| --- | --- |
| Plants | 279 |
| Terrestrial Mammals | 261 |
| Aquatic Mammals | 192 |
| Birds | 300 |
| Invertebrates | 417 |
| Fish | 333 |
| Herpetofauna | 253 |
| *Amphibians* | *215* |
| *Reptiles* | *135* |
| Fungi | 215 |
| Microorganisms | 194 |
| **Total** | **2444** |

#### 5.1.2 Systematic search

The following information details the final, systematically determined search strings for each taxa.

##### Plants

Scopus: (plant OR plants) AND (biodiversity OR diversity OR species) AND survey* AND (ecosystem OR environment). WOS: (plant OR plants) AND (biodiversity OR diversity OR species) AND survey* AND ecosystem.

##### Birds

Scopus and WOS: (bird OR birds) AND (biodiversity OR diversity OR species) AND (survey OR surveys) AND habitat.

##### Mammal

Scopus and WOS: (mammal OR mammals) AND (biodiversity OR diversity OR species) AND (survey OR surveys).

##### Marine Mammal

Scopus and WOS: (marine mammal OR marine mammals) AND survey*.

##### Invertebrate

Scopus and WOS: (invertebrate OR invertebrates) AND (biodiversity OR diversity OR species) AND (survey OR surveys).

##### Herpetofauna

Scopus and WOS: (amphibia OR amphibian OR amphibians OR reptile OR reptiles OR anuran*) AND (biodiversity OR diversity) AND (survey OR surveys).

##### Fish

Scopus and WOS: (fish OR fishes) AND (biodiversity OR diversity OR species) AND survey*.

##### Fungi

Scopus and WOS: (fungi OR fungal) AND (biodiversity OR diversity OR species) AND survey*.

##### Microorganisms

Scopus and WOS: (bacteria OR bacterial OR rna OR microbal OR microbe) AND (biodiversity OR diversity OR communities) AND survey*.

### 5.2 Data Extraction

Our team reviewed all suitable papers and extracted relevant information into spreadsheets; for each paper we extracted DOI, taxa, publication date, spatial information, biome, methods and sub methods. In order to ensure consistency among reviewers, we provided a starter training session, notes to refer to, and data collection sheets were drop down boxes where possible. One person checked through all data collection sheets and resolved any queries or anomalies in data recorded.

### 5.3 Data Analysis

A high throughput method of measuring diversity is one that can efficiently collect or process a lot of data in a short amount of time. As new technologies are emerging it is expected that their use across biodiversity studies will increase overtime. We tested how the use of higher throughput methods has changed overtime, versus low throughput methods. To do this we used expert opinion to determine if each of the 14 methods were high or low throughput by considering how efficiently and effectively they can collect information across different spatial and temporal scales. eDNA, acoustics, camera trapping, auonomous vehicles and remote sensing are considered to be high throughout methods and the remaining methods are considered to be low throughout. Table 2.

**Table 2.** Table showing each of the 14 methodologies and whether they are high or low throughput.

| Method | Throughput |
| --- | --- |
| Acoustics | High |
| Camera Trap | High |
| eDNA | High |
| Light Trap | Low |
| Mammal Trap | Low |
| Marine Transect | Low |
| Pitfall Trap | Low |
| Point Count | Low |
| Quadrat | Low |
| Remote Sensing | High |
| SCUBA Survey | Low |
| Terrestrial Transect | Low |
| Unmanned Vehicle | High |
| Visual Encounters | Low |

All analyses were performed using R Statistical Software (v4.1; (**tidyv**; **dplyr**)). Once the data had been collated raw data were subset by taxa and methodology to perform descriptive statistics using packages `dplyr` and `tidyverse` (**tidyv**; **dplyr**). To understand richness and evenness of taxa represented across methods we used package `vegan` (**vegan**) to quantify simple richness and Simpson’s Evenness Index (*D*), where a score of 1 indicates low evenness of taxa or methods. To test if the rate of change in published biodiversity studies were higher than general scientific studies, we used a dataset of yearly scientific publications globally (World Bank 2024), which is calculated from scraping studies from publication sites on the web, as we did in this study. This data were then used to calculate annual and cumulative rate of change from 2000-2020, as published studies were not yet available after 2020. Total percent change for general scientific studies from 2000:2022 was just 188%, a 10-fold decrease when compared to biodiversity studies.

To analyse temporal trends in the publication of studies overtime we used polynomial poisson regression with a quadratic term for year due to the non-linear, curved nature of our data and to model count data. To analyse temporal trends in taxa and methods we used binomial models to assess changes in proportions of different groups given changes in overall sampling, with number of studies per taxa or method and total number of studies combined as the response variable and year as the explanatory variable in package `stats` (**r·core·team·r·2024**). If the relationship was non linear, determined through assessing residual plots and visualising trends in the data, we used polynomial regression with a quadratic term for year. Polynomial models were chosen as they offer increased flexibility in modeling non-linear relationships and allowed us to capture curvatures in data that linear models cannot, providing a better fit and potentially more accurate predictions. Quadratic terms were applied where data were not linear, being careful not to overfit with additional terms. To determine if the polynomial model was a better fit we assessed residual plots and the *p* value of the polynomial term, with a *p <* 0.05 indicating that there is a curve-linear relationship. We also determined if inclusion of the quadratic term significantly improved model fit across all models when compared to a simple linear regression model and improved model diagnostics. To determine final model fit we created a null model and used the anova function in package `stats` (**r·core·team·r·2024**) to compare that and the full model. We used *p, χ*2 extracted from the model and pseudo *r*^2^ values to determine model significance and strength. Since a true *r*^2^ doesn’t exist in logistic regression pseudo *r*^2^ were used to assesses the goodness-of-fit of the model using package `DescTools` (**signorell·a··desctools·2024**). pseudo *r*^2^ values serve as approximations to quantify the model’s predictive power and the amount of variance explained by the independent variables. To test the significance of model terms, devience was calculated by comparing the residual deviance of the null and final model.

To understand if the distribution of studies across biomes and continents deviated from expected values, which we assumed to be equal, we used the *χ*2 goodness-of-fit test, chosen as it is used to determine if a sample of data is likely to have come from a specific theoretical distribution. All plots were generated using the package `ggplot2` (**ggplot2**).

## 6 Acknowledgements

This work was funded by BP. Additionally, the Pearse Lab is supported by UK Research and Innovation (UKRI) grants BB/Y008766/1, NE/X00547X/1, and NE/X013022/1, as well as by the Alan Turing Institute and the Singapore Green Finance Centre (Nature-Related Financial Risks in Southeast Asia). The authors gratefully acknowledge this support, which made this research possible.

## References

Allan, Blake M. et al. (2018). “Futurecasting ecological research: the rise of technoecology”. en. In: Ecosphere 9.5. eprint: https://onlinelibrary.wiley.com/doi/pdf/10.1002/ecs2.2163, pe02163. issn: 2150-8925. doi: 10.1002/ecs2.2163. url: https://onlinelibrary.wiley.com/doi/abs/10.1002/ecs2.2163 (visited on 10/16/2023).

Almond, R.E.A. et al. (2022). Living Planet Report 2022 – Building a naturepositive society.

Beng, Kingsly C. and Richard T. Corlett (June 2020). “Applications of environmental DNA (eDNA) in ecology and conservation: opportunities, challenges and prospects”. en. In: Biodiversity and Conservation 29.7, pp. 2089–2121. issn: 1572-9710. doi: 10.1007/s10531-020-01980-0. url: https://doi.org/10.1007/s10531-020-01980-0 (visited on 10/13/2023).

Borja, Angel et al. (July 2024). “Innovative and practical tools for monitoring and assessing biodiversity status and impacts of multiple human pressures in marine systems”. In: Environmental Monitoring and Assessment 196.8, p. 694. issn: 1573-2959. doi: 10.1007/s10661-024-12861-2. url: https://doi.org/10.1007/s10661-024-12861-2.

Boyd, Ian L., W. Don Bowen, and Sara J. Iverson (Aug. 2010). Marine Mammal Ecology and Con-servation: A Handbook of Techniques. en. Google-Books-ID: QdkTDAAAQBAJ. Oxford University Press. isbn: 978-0-19-921656-7.

CBD (Oct. 2010). Decision X/2, The Strategic Plan for Biodiversity 2011–2020 and the Aichi Biodiversity Targets. (visited on 07/31/2024).

CBD (2022). Decision 15/5: Monitoring framework for the Kunming-Montreal Global Biodiversity Framework. https://www.cbd.int/doc/decisions/cop-15/cop-15-dec-05-en.pdf. Accessed: 2025-10-11.

Chowdhury, Shawan et al. (June 2023). “Increasing biodiversity knowledge through social media: A case study from tropical Bangladesh”. In: BioScience 73.6, pp. 453–459. issn: 0006-3568. doi: 10.1093/biosci/biad042. url: https://doi.org/10.1093/biosci/biad042 (visited on 03/11/2024).

Clark, J. Alan and Robert M. May (July 2002). “Taxonomic Bias in Conservation Research”. In: Science 297.5579. Publisher: American Association for the Advancement of Science, pp. 191–192. doi: 10.1126/science.297.5579.191b. url: https://www.science.org/doi/10.1126/science.297.5579.191b (visited on 09/22/2023).

Crunchant, Anne-Sophie et al. (2020). “Listening and watching: Do camera traps or acoustic sensors more efficiently detect wild chimpanzees in an open habitat?” en. In: Methods in Ecology and Evolution 11.4. eprint: https://onlinelibrary.wiley.com/doi/pdf/10.1111/2041-210X.13362, xpp. 542–552. issn: 2041-210X. doi: 10.1111/2041-210X.13362. url: https://onlinelibrary.wiley.com/doi/abs/10.1111/2041-210X.13362 (visited on 09/15/2023).

Delisle, Zackary J. et al. (2021). “Next-Generation Camera Trapping: Systematic Review of Historic Trends Suggests Keys to Expanded Research Applications in Ecology and Conservation”. In: Frontiers in Ecology and Evolution 9. issn: 2296-701X. url: https://www.frontiersin.org/articles/10.3389/fevo.2021.617996 (visited on 10/13/2023).

Di Marco, Moreno et al. (Apr. 2017). “Changing trends and persisting biases in three decades of conservation science”. In: Global Ecology and Conservation 10, pp. 32–42. issn: 2351-9894. doi:10.1016/j.gecco.2017.01.008. url: https://www.sciencedirect.com/science/article/pii/S2351989417300148 (visited on 09/28/2023).

Doan, Tiffany M. (2003). “Which Methods Are Most Effective for Surveying Rain Forest Herpetofauna?” In: Journal of Herpetology 37.1. Publisher: Society for the Study of Amphibians and Reptiles, pp. 72–81. issn: 0022-1511. url: https://www.jstor.org/stable/1565833 (visited on 09/15/2023).

Enari, Hiroto et al. (Mar. 2019). “An evaluation of the efficiency of passive acoustic monitoring in detecting deer and primates in comparison with camera traps”. In: Ecological Indicators 98, pp. 753–762. issn: 1470-160X. doi: 10.1016/j.ecolind.2018.11.062. url: https://www.sciencedirect.com/science/article/pii/S1470160X18309257 (visited on 09/15/2023).

European Commission (2014). Study on specific design elements of biodiversity offsets – Biodiversity metrics & mechanisms for securing long term conservation benefits: 2nd edition, final report. en. Tech. rep. Publications Office. url: https://data.europa.eu/doi/10.2779/010692 (visited on 07/26/2024).

Farooq, Harith et al. (May 2021). “Mapping Africa’s Biodiversity: More of the Same Is Just Not Good Enough”. In: Systematic Biology 70.3, pp. 623–633. issn: 1063-5157. doi: 10.1093/sysbio/syaa090. url: https://doi.org/10.1093/sysbio/syaa090 (visited on 03/17/2024).

Ficetola, Gentile Francesco et al. (Aug. 2008). “Species detection using environmental DNA from water samples”. en. In: Biology Letters 4.4, pp. 423–425. issn: 1744-9561, 1744-957X. doi: 10.1098/rsbl.2008.0118. url: https://royalsocietypublishing.org/doi/10.1098/rsbl.2008.0118 (visited on 07/28/2024).

GBIF (2024). GBIF. url: https://www.gbif.org/ (visited on 07/26/2024).

Gill, M et al. (2017). A Sourcebook of Methods and Procedures for Monitoring Essential Biodiversity Variables in Tropical Forests with Remote Sensing., The Netherlands: GOFC-GOLD Land Cover Project Office. isbn: 2542-6729.

Goffredo, Stefano, Francesco Pensa, et al. (Dec. 2010). “Unite research with what citizens do for fun: “recreational monitoring” of marine biodiversity”. en. In: Ecological Applications 20.8, pp. 2170– 2187. issn: 1051-0761, 1939-5582. doi: 10.1890/09-1546.1. url: https://esajournals.onlinelibrary.wiley.com/doi/10.1890/09-1546.1 (visited on 07/28/2024).

Goffredo, Stefano, Corrado Piccinetti, and Francesco Zaccanti (Dec. 2004). “Volunteers in Marine Conservation Monitoring: a Study of the Distribution of Seahorses Carried Out in Collaboration with Recreational Scuba DiversVoluntarios en el Monitoreo de Conservación Marina: un Estudio de Distribución de Caballitos de Mar Llevado a Cabo con Buzos Scuba Recreativos”. en. In: Conservation Biology 18.6. Publisher: John Wiley & Sons, Ltd, pp. 1492–1503. issn: 1523-1739. doi:10.1111/j.1523-1739.2004.00015.x. url: https://conbio.onlinelibrary.wiley.com/doi/10.1111/j.1523-1739.2004.00015.x (visited on 07/28/2024).

Gooday, Oliver Jordan et al. (June 2018). “An assessment of thermal-image acquisition with an unmanned aerial vehicle (UAV) for direct counts of coastal marine mammals ashore”. In: Journal of Unmanned Vehicle Systems 6.2. Publisher: NRC Research Press, pp. 100–108. issn: 2291-3467. doi: 10.1139/juvs-2016-0029. url: https://cdnsciencepub.com/doi/10.1139/juvs-2016-0029 (visited on 02/17/2025).

Hallmann, Caspar A. et al. (Oct. 2017). “More than 75 percent decline over 27 years in total flying insect biomass in protected areas”. en. In: PLOS ONE 12.10. Publisher: Public Library of Science, e0185809. issn: 1932-6203. doi: 10.1371/journal.pone.0185809. url: https://journals.plos.org/plosone/article?id=10.1371/journal.pone.0185809 (visited on 07/31/2024).

Hermoso, Maibe, Soledad Narváez, and Martin Thiel (Feb. 2021). “Engaging recreational scuba divers in marine citizen science: Differences according to popularity of the diving area”. en. In: Aquatic Conservation: Marine and Freshwater Ecosystems 31.2. Publisher: John Wiley & Sons, Ltd, pp. 441–455. issn: 1099-0755. doi: 10.1002/aqc.3466. url: https://onlinelibrary.wiley.com/doi/10.1002/aqc.3466 (visited on 07/28/2024).

Hoffmann, Anke et al. (2010). Field Methods and Techniques for Monitoring Mammals. ldots recording techniques.

Hughes, Terry P. et al. (Jan. 2018). “Spatial and temporal patterns of mass bleaching of corals in the Anthropocene”. In: Science 359.6371. Publisher: American Association for the Advancement of Science, pp. 80–83. doi: 10.1126/science.aan8048. url: https://www.science.org/doi/10.1126/science.aan8048 (visited on 02/17/2025).

iNaturalist (2024). iNaturalist. en-US. url: https://www.inaturalist.org/ (visited on 07/26/2024).

IPBES (May 2019). Global assessment report on biodiversity and ecosystem services of the Intergovernmental Science-Policy Platform on Biodiversity and Ecosystem Services. eng. Tech. rep. Zenodo. doi: 10.5281/zenodo.6417333. url: https://zenodo.org/records/6417333 (visited on 03/18/2024).

Junker, Jessica et al. (Sept. 2020). “A Severe Lack of Evidence Limits Effective Conservation of the World’s Primates”. en. In: BioScience 70.9, pp. 794–803. issn: 0006-3568, 1525-3244. doi: 10.1093/biosci/biaa082. url: https://academic.oup.com/bioscience/article/70/9/794/5896003 (visited on 07/26/2024).

Kelaher, Brendan P. et al. (Jan. 2019). “Assessing variation in assemblages of large marine fauna off ocean beaches using drones”. en. In: Marine and Freshwater Research 71.1. Publisher: CSIRO PUBLISHING, pp. 68–77. issn: 1448-6059. doi: 10.1071/MF18375. url: https://www.publish.csiro.au/mf/MF18375 (visited on 02/17/2025).

Larigauderie, Anne and Harold A Mooney (May 2010). “The Intergovernmental science-policy Platform on Biodiversity and Ecosystem Services: moving a step closer to an IPCC-like mechanism for biodiversity”. In: Current Opinion in Environmental Sustainability 2.1, pp. 9–14. issn: 1877-3435. doi: 10.1016/j.cosust.2010.02.006. url: https://www.sciencedirect.com/science/article/pii/S1877343510000072 (visited on 01/15/2025).

Leung, Brian et al. (Dec. 2020). “Clustered versus catastrophic global vertebrate declines”. en. In: Nature 588.7837. Publisher: Nature Publishing Group, pp. 267–271. issn: 1476-4687. doi: 10.1038/s41586-020-2920-6. url: https://www.nature.com/articles/s41586-020-2920-6 (visited on 01/13/2025).

Linke, Simon et al. (Mar. 2018). “Freshwater ecoacoustics as a tool for continuous ecosystem monitoring”. In: Frontiers in Ecology and the Environment 16. doi: 10.1002/fee.1779.

Lister, Bradford C. and Andres Garcia (Oct. 2018). “Climate-driven declines in arthropod abundance restructure a rainforest food web”. In: Proceedings of the National Academy of Sciences 115.44. Publisher: Proceedings of the National Academy of Sciences, E10397–E10406. doi: 10.1073/pnas.1722477115. url: https://www.pnas.org/doi/full/10.1073/pnas.1722477115 (visited on 07/31/2024).

Lynggaard, Christina et al. (2022). “Airborne environmental DNA for terrestrial vertebrate community monitoring”. In: Current Biology 32.3, 701–707.e5. issn: 0960-9822. doi: 10.1016/j.cub.2021.12.014. url: https://www.sciencedirect.com/science/article/pii/S0960982221016900.

Maasri, Alain et al. (2022). “A global agenda for advancing freshwater biodiversity research”. en. In: Ecology Letters 25.2. eprint: https://onlinelibrary.wiley.com/doi/pdf/10.1111/ele.13931, xpp. 255– 263. issn: 1461-0248. doi: 10.1111/ele.13931. url: https://onlinelibrary.wiley.com/doi/abs/10.1111/ele.13931 (visited on 09/28/2023).

MacWilliams, Ryley, Seojin Kim, and Rebecca J. Trueman (Oct. 2024). “Bugs and bots: how technology is changing the game in biodiversity monitoring”. In: Biodiversity 25.4. Publisher: Taylor & Francis, pp. 295–296. issn: 1488-8386. doi: 10.1080/14888386.2024.2419829. url: https://doi.org/10.1080/14888386.2024.2419829.

Mallet, Delphine and Dominique Pelletier (June 2014). “Underwater video techniques for observing coastal marine biodiversity: A review of sixty years of publications (1952–2012)”. In: Fisheries Research 154, pp. 44–62. issn: 0165-7836. doi: 10.1016/j.fishres.2014.01.019. url: https://www.sciencedirect.com/science/article/pii/S0165783614000356 (visited on 02/16/2025).

Marshall, Erica et al. (Jan. 2020). “What are we measuring? A review of metrics used to describe biodiversity in offsets exchanges”. In: Biological Conservation 241, p. 108250. issn: 0006-3207. doi: 10.1016/j.biocon.2019.108250. url: https://www.sciencedirect.com/science/article/pii/S0006320719309620 (visited on 07/26/2024).

Marvin, David C. et al. (July 2016). “Integrating technologies for scalable ecology and conservation”. In: Global Ecology and Conservation 7, pp. 262–275. issn: 2351-9894. doi: 10.1016/j.gecco.2016.07.002. url: https://www.sciencedirect.com/science/article/pii/S2351989416300592 (visited on 10/16/2023).

McGeady, Ryan et al. (July 2023). “A review of new and existing non-extractive techniques for monitoring marine protected areas”. English. In: Frontiers in Marine Science 10. Publisher: Frontiers. issn: 2296-7745. doi: 10.3389/fmars.2023.1126301. url: https://www.frontiersin.org/journals/marine-science/articles/10.3389/fmars.2023.1126301/full (visited on 02/17/2025).

Meinam, Martina et al. (2023). “Importance of fish biodiversity conservation and management”. en. In: International Journal of Science and Research Archive 9.2. Number: 2 Publisher: International Journal of Science and Research Archive, pp. 387–391. issn: 2582-8185, 2582-8185. doi: 10.30574/ijsra.2023.9.2.0513. url: https://ijsra.net/content/importance-fish-biodiversity-conservation-and-management (visited on 07/31/2024).

Navarro, Laetitia M et al. (Dec. 2017). “Monitoring biodiversity change through effective global coordination”. In: Current Opinion in Environmental Sustainability 29, pp. 158–169. issn: 1877-3435. doi: 10.1016/j.cosust.2018.02.005. url: https://www.sciencedirect.com/science/article/pii/S1877343517301665 (visited on 01/15/2025).

Newbold, Tim et al. (Apr. 2015). “Global effects of land use on local terrestrial biodiversity”. en. In: Nature 520.7545. Publisher: Nature Publishing Group, pp. 45–50. issn: 1476-4687. doi: 10.1038/nature14324. url: https://www.nature.com/articles/nature14324 (visited on 07/26/2024).

Nisa, Zahidah Afrin, Clive Schofield, and Francis C. Neat (Feb. 2022). “Work Below Water: The role of scuba industry in realising sustainable development goals in small island developing states”. In: Marine Policy 136, p. 104918. issn: 0308-597X. doi: 10.1016/j.marpol.2021.104918. url: https://www.sciencedirect.com/science/article/pii/S0308597X21005297 (visited on 07/28/2024).

Oliver, Ruth Y. et al. (May 2023). “Camera trapping expands the view into global biodiversity and its change”. In: Philosophical Transactions of the Royal Society B: Biological Sciences 378.1881. Publisher: Royal Society, p. 20220232. doi: 10.1098/rstb.2022.0232. url: https://royalsocietypublishing.org/doi/10.1098/rstb.2022.0232 (visited on 03/11/2024).

Pearce, J.L. et al. (Apr. 2005). “Pitfall trap designs to maximize invertebrate captures and minimize captures of nontarget vertebrates”. en. In: The Canadian Entomologist 137.2, pp. 233–250. issn: 0008-347X, 1918-3240. doi: 10.4039/n04-029. url: https://www.cambridge.org/core/product/identifier/S0008347X00002431/type/journal_article (visited on 03/16/2024).

Pereira, H. M. et al. (Jan. 2013). “Essential Biodiversity Variables”. In: Science 339.6117. Publisher: American Association for the Advancement of Science, pp. 277–278. doi: 10.1126/science.1229931. url: https://www.science.org/doi/10.1126/science.1229931 (visited on 01/15/2025).

Pimm, Stuart L. et al. (Nov. 2015). “Emerging Technologies to Conserve Biodiversity”. en. In: Trends in Ecology & Evolution 30.11, pp. 685–696. issn: 01695347. doi: 10.1016/j.tree.2015.08.008. url: https://linkinghub.elsevier.com/retrieve/pii/S0169534715002128 (visited on 10/27/2023).

Pistón, N et al. (2023). GEO BON Working Group on Ecosystem Services: Implementation Plan 2023-2026. Tech. rep. url: https://geobon.org/ebvs/ecosystem-services/.

Purvis, Andy and Andy Hector (May 2000). “Getting the measure of biodiversity”. en. In: Nature 405.6783. Publisher: Nature Publishing Group, pp. 212–219. issn: 1476-4687. doi: 10.1038/35012221. url: https://www.nature.com/articles/35012221 (visited on 07/26/2024).

Rasmussen, Jeppe H., Dan Stowell, and Elodie F. Briefer (2024). “Sound evidence for biodiversity monitoring”. In: Science 385.6705. eprint: https://www.science.org/doi/pdf/10.1126/science.adh2716, xpp. 138–140. doi: 10.1126/science.adh2716. url: https://www.science.org/doi/abs/10.1126/science.adh2716.

Reynolds, Sam A. et al. (Feb. 2025). “The potential for AI to revolutionize conservation: a horizon scan”. In: Trends in Ecology & Evolution 40.2, pp. 191–207. issn: 0169-5347. doi: 10.1016/j.tree.2024.11.013. url: https://www.sciencedirect.com/science/article/pii/S0169534724002866 (visited on 02/16/2025).

Roy, D.B. et al. (May 2024). “Towards a standardized framework for AI-assisted, image-based monitoring of nocturnal insects”. In: Philosophical Transactions of the Royal Society B 379. doi: 10.1098/rstb.2023.0108.

Ruppert, Krista M., Richard J. Kline, and Md Saydur Rahman (Jan. 2019). “Past, present, and future perspectives of environmental DNA (eDNA) metabarcoding: A systematic review in methods, monitoring, and applications of global eDNA”. In: Global Ecology and Conservation 17, e00547. issn: 2351-9894. doi: 10.1016/j.gecco.2019.e00547. url: https://www.sciencedirect.com/science/article/pii/S2351989418303500 (visited on 03/11/2024).

Santini, Luca et al. (Sept. 2017). “Assessing the suitability of diversity metrics to detect biodiversity change”. In: Biological Conservation. SI:Measures of biodiversity 213, pp. 341–350. issn: 0006-3207. doi: 10.1016/j.biocon.2016.08.024. url: https://www.sciencedirect.com/science/article/pii/S0006320716303305 (visited on 07/26/2024).

Sayer, Catherine A. et al. (Jan. 2025). “One-quarter of freshwater fauna threatened with extinction”. en. In: Nature. Publisher: Nature Publishing Group, pp. 1–8. issn: 1476-4687. doi: 10.1038/s41586-024-08375-z. url: https://www.nature.com/articles/s41586-024-08375-z (visited on 01/13/2025).

Scholes, Robert J. et al. (2012). “Building a global observing system for biodiversity”. In: Current Opinion in Environmental Sustainability 4.1, pp. 139–146. issn: 1877-3435. doi: 10.1016/j.cosust.2011.12.005. url: https://www.sciencedirect.com/science/article/pii/S1877343511001394.

Speaker, Talia et al. (2022). “A global community-sourced assessment of the state of conservation technology”. en. In: Conservation Biology 36.3. eprint: https://onlinelibrary.wiley.com/doi/pdf/10.1111/cobi.13871, e13871. issn: 1523-1739. doi: 10.1111/cobi.13871. url: https://onlinelibrary.wiley.com/doi/abs/10.1111/cobi.13871 (visited on 07/31/2024).

Stephenson, P. J. (2020). “Technological advances in biodiversity monitoring: applicability, opportunities and challenges”. In: Current Opinion in Environmental Sustainability 45, pp. 36–41. issn: 1877-3435. doi: 10.1016/j.cosust.2020.08.005. url: https://www.sciencedirect.com/science/article/pii/S1877343520300592.

Stephenson, P. J., Yaa Ntiamoa-Baidu, and John P. Simaika (2020). “The Use of Traditional and Modern Tools for Monitoring Wetlands Biodiversity in Africa: Challenges and Opportunities”. In: Frontiers in Environmental Science 8. issn: 2296-665X. doi: 10.3389/fenvs.2020.00061. url: https://www.frontiersin.org/journals/environmental-science/articles/10.3389/fenvs.2020.00061.

Thomas, Chris D., T. Hefin Jones, and Sue E. Hartley (2019). ““Insectageddon”: A call for more robust data and rigorous analyses”. en. In: Global Change Biology 25.6. eprint: https://onlinelibrary.wiley.com/doi/pdf/10.111 pp. 1891–1892. issn: 1365-2486. doi: 10.1111/gcb.14608. url: https://onlinelibrary.wiley.com/doi/abs/10.1111/gcb.14608 (visited on 07/31/2024).

Trimble, Morgan J. and Rudi J. van Aarde (2012). “Geographical and taxonomic biases in research on biodiversity in human-modified landscapes”. en. In: Ecosphere 3.12. eprint: https://onlinelibrary.wiley.com/doi/pdf/1000299.1, art119. issn: 2150-8925. doi: 10.1890/ES12-00299.1. url: https://onlinelibrary.wiley.com/doi/abs/10.1890/ES12-00299.1 (visited on 09/22/2023).

Troudet, Julien et al. (Aug. 2017). “Taxonomic bias in biodiversity data and societal preferences”. en. In: Scientific Reports 7.1. Number: 1 Publisher: Nature Publishing Group, p. 9132. issn: 2045-2322. doi: 10.1038/s41598-017-09084-6. url: https://www.nature.com/articles/s41598-017-09084-6 (visited on 09/22/2023).

Vèlez, Juliana et al. (2023). “An evaluation of platforms for processing camera-trap data using artificial intelligence”. In: Methods in Ecology and Evolution 14.2, pp. 459–477. doi: 10.1111/2041-210X.14044. eprint: https://besjournals.onlinelibrary.wiley.com/doi/pdf/10.1111/2041-210X.14044. url: https://besjournals.onlinelibrary.wiley.com/doi/abs/10.1111/2041-210X.14044.

Vellend, Mark et al. (Nov. 2013). “Global meta-analysis reveals no net change in local-scale plant biodiversity over time”. In: Proceedings of the National Academy of Sciences 110.48. Publisher: Proceedings of the National Academy of Sciences, pp. 19456–19459. doi: 10.1073/pnas.1312779110. url: https://www.pnas.org/doi/full/10.1073/pnas.1312779110 (visited on 07/26/2024).

Venter, J. Craig et al. (Apr. 2004). “Environmental Genome Shotgun Sequencing of the Sargasso Sea”. In: Science 304.5667. Publisher: American Association for the Advancement of Science, pp. 66–74. doi: 10.1126/science.1093857. url: https://www.science.org/doi/10.1126/science.1093857 (visited on 07/28/2024).

Wägele J. Wolfgang et al. (Mar. 2022). “Towards a multisensor station for automated biodiversity monitoring”. In: Basic and Applied Ecology 59, pp. 105–138. issn: 1439-1791. doi: 10.1016/j.baae.2022.01.003. url: https://www.sciencedirect.com/science/article/pii/S1439179122000032 (visited on 02/17/2025).

Whitworth, Andrew et al. (Jan. 2017). “Methods matter: Different biodiversity survey methodologies identify contrasting biodiversity patterns in a human modified rainforest — A case study with amphibians”. In: Ecological Indicators 72, pp. 821–832. doi: 10.1016/j.ecolind.2016.08.055.

World Bank (2024). Annual articles published in scientific and technical journals-Our World in Data. url: https://ourworldindata.org/grapher/scientific-and-technical-journal-articles%20%5Bonline%20resource%5D.

Zhao, Zi-Hua et al. (2013). “Solving the pitfalls of pitfall trapping: a two-circle method for density estimation of ground-dwelling arthropods”. en. In: Methods in Ecology and Evolution 4.9. eprint: https://onlinelibrary.wiley.com/doi/pdf/10.1111/2041-210X.12083, xpp. 865–871. issn: 2041-210X. doi: 10.1111/2041-210X.12083. url: https://onlinelibrary.wiley.com/doi/abs/10.1111/2041-210X.12083 (visited on 03/16/2024).

