## Supplementary material for "Systematic review of biodiversity monitoring shows gradual rise of scaleable and automated methods and inherent spatial and taxonomic biases"

### 1 Diversity of methods and taxa

| Method | Richness of taxa | Simpsons evenness Index (*D*) |
| --- | --- | --- |
| Quadrat | 8 | 0.19097267 |
| Visual Encounter | 8 | 0.213072195 |
| eDNA | 9 | 0.2246872 |
| Point Count | 9 | 0.238085867 |
| Camera Trap | 6 | 0.263815124 |
| Marine Transect | 7 | 0.269578892 |
| Unmanned Vehicle | 7 | 0.285184049 |
| Acoustics | 6 | 0.347915463 |
| Terrestrial Transect | 6 | 0.357355051 |
| SCUBA | 4 | 0.447047376 |
| Remote Sensing | 2 | 0.547471851 |
| Pitfall trap | 3 | 0.588846813 |
| Light Trap | 1 | 1 |
| Mammal Trap | 1 | 1 |

Table 1: **Richness and evenness of taxa represented across methods.** This table shows the Richness of taxa, which represents the number of taxonomic groups found within each method and evenness of taxa, which shows how evenly spread taxonomic studies are within a method.

| Taxa | Richness of methods | Simpsons evenness Index (*D*) |
| --- | --- | --- |
| Plant | 10 | 0.16022437 |
| Herpetofauna | 10 | 0.179660388 |
| Invertebrate | 12 | 0.194094052 |
| Terrestrial.Mammal | 11 | 0.19417287 |
| Aquatic.Mammal | 8 | 0.239610924 |
| Birds | 9 | 0.242056514 |
| Fish | 9 | 0.253792763 |
| Fungi | 5 | 0.328935521 |
| Microorganism | 3 | 0.352828911 |

Table 2: **Richness and evenness of methods represented within a taxa.** This table shows richness of methods, which represents the number of methods used to study each taxa and evenness of methods, which shows how evenly spread methods are across within a taxonomic group.

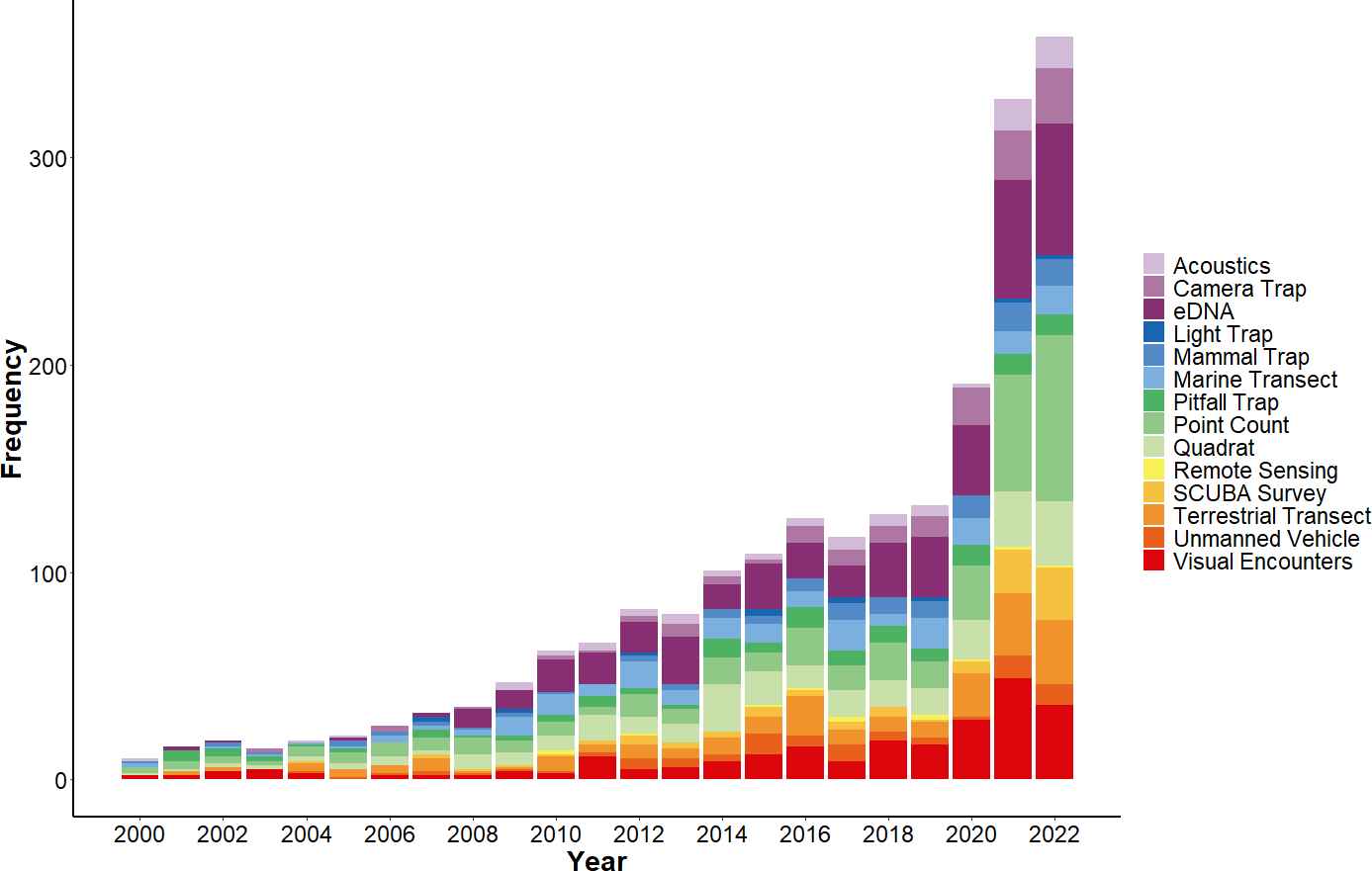

Figure 2: **Frequency of published studies from 2000–2022 per methodology.** The horizontal axis displays the year 2000:2022 in ascending order. The vertical axis shows the frequency of studies each year. Each stacked section and of the bar represents an individual methodology with representa- tive colours detailed in the legend. The figure highlights the rapid increase in the frequency of studies from 2020 and the change in number of studies over time per method

### 2 Taxonomic Figures

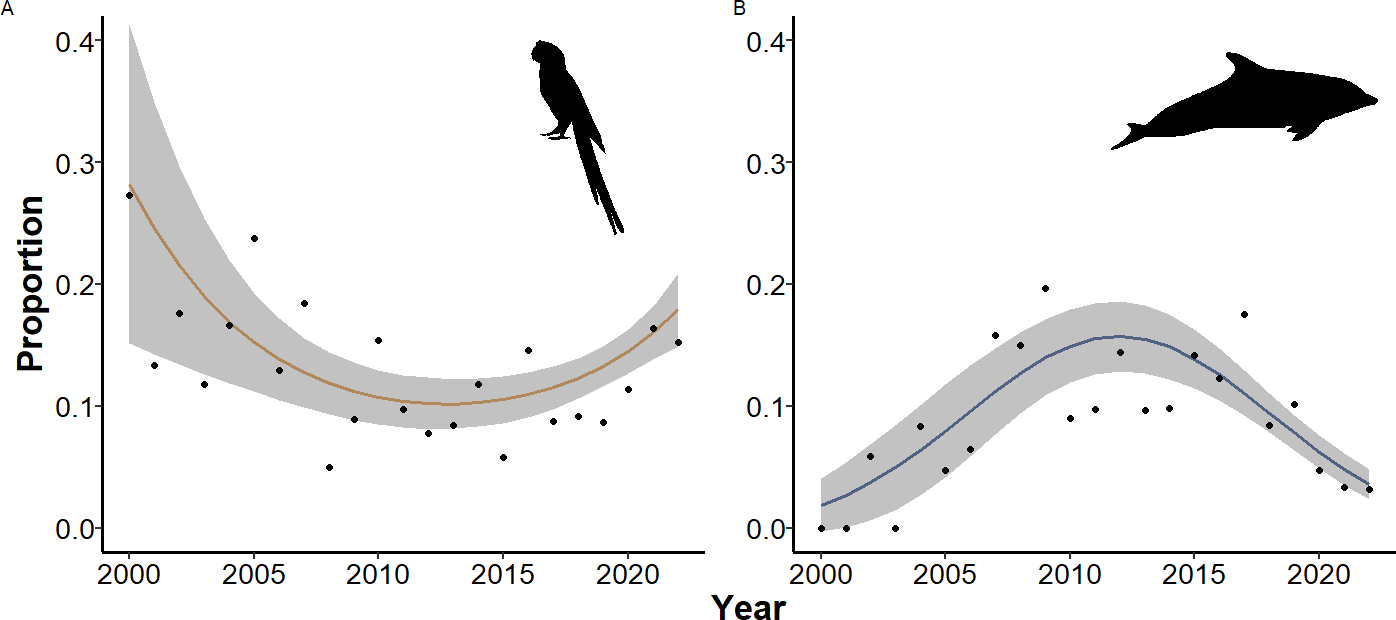

Figure 1: **Proportional change in published studies for birds and marine mammals over- time.** The horizontal axis displays years from 2000–2022.The vertical axis shows the proportional change in the number of studies for each taxonomic group, with year on the horizontal axis. Of the nine taxonomic groups included in the meta-analysis we display here only the significant results for, A. birds and B. Marine mammals. Regression with a quadratic term for year showed a significant relationship between time and proportion of studies on birds (*p <* 0*.*001, *χ*2 = 17*.*6_(22_*_,_*_20)_, pseudo- *r*^2^ = 0*.*88) and aquatic mammals after 2010 (*p <* 0*.*001, *χ*2 = 52*.*91_(22_*_,_*_20)_, pseudo-*r*^2^ = 0*.*88).

3 **Frequency change in study model outputs**

|  | Estimate | Std. Error | z value | Pr(*>\|*z*\|*) |
| --- | --- | --- | --- | --- |
| (Intercept) | 4.1863 | 0.0312 | 134.27 | 0.0000 |
| poly(year, 2)1 | 4.5198 | 0.1604 | 28.18 | 0.0000 |
| poly(year, 2)2 | 0.4199 | 0.1209 | 3.47 | 0.0005 |

Table 3: **Model Output: Change in frequency of studies over time.** Poisson regression with a quadratic term for year.

### 4 Taxonomic model outputs

| Estimate | Std. Error | z value | Pr(*>\|*z*\|*) |
| --- | --- | --- | --- |
| (Intercept) 87.4370 | 19.2859 | 4.53 | 0.0000 |
| year -0.0442 | 0.0096 | -4.62 | 0.0000 |

Table 4: **Model Output: Change in proportion of invertebrate studies overtime.** Binomial regression shows a significant relationship between the proportion of studies and year.

|  | Estimate | Std. Error | z value | Pr(*>\|*z*\|*) |
| --- | --- | --- | --- | --- |
| (Intercept) | -1.8169 | 0.0923 | -19.69 | 0.0000 |
| poly(year, 2)1 | -0.8509 | 0.4579 | -1.86 | 0.0631 |
| poly(year, 2)2 | 1.4380 | 0.3500 | 4.11 | 0.0000 |

Table 5: **Model Output:proportion of bird overtime.** Binomial regression with a quadratic term for year shows a significant curvelinear relationship between the proportion of studies and year.

|  | Estimate | Std. Error | z value | Pr(*>\|*z*\|*) |
| --- | --- | --- | --- | --- |
| (Intercept) | -2.3924 | 0.1389 | -17.22 | 0.0000 |
| poly(year, 2)1 | 0.9648 | 0.7921 | 1.22 | 0.2232 |
| poly(year, 2)2 | -2.9933 | 0.5561 | -5.38 | 0.0000 |

Table 6: **Model Output:proportion of aquatic mammal overtime.** Binomial regression with a quadratic term for year shows a significant curvelinear relationship between the proportion of studies and year.

|  | Estimate | Std. Error | z value | Pr(*>\|*z*\|*) |
| --- | --- | --- | --- | --- |
| (Intercept) | -2.7613 | 0.1456 | -18.97 | 0.0000 |
| poly(year, 2)1 | 1.3020 | 0.6228 | 2.09 | 0.0366 |
| poly(year, 2)2 | 4.3020 | 0.4526 | 9.51 | 0.0000 |

Table 7: **Model Output:proportion of fish studies overtime.** Binomial regression with a quadratic term for year shows a significant curvelinear relationship between the proportion of studies and year.

|  | Estimate | Std. Error | z value | Pr(*>\|*z*\|*) |
| --- | --- | --- | --- | --- |
| (Intercept) | -10.7766 | 27.9823 | -0.39 | 0.7001 |
| year | 0.0043 | 0.0139 | 0.31 | 0.7593 |

Table 8: **Model Output:proportion of fungi studies overtime.** Binomial regression shows no significant relationship between the proportion of studies and year.

|  | Estimate | Std. Error | t value | Pr(*>\|*t*\|*) |
| --- | --- | --- | --- | --- |
| (Intercept) | -54.4778 | 31.6439 | -1.72 | 0.0998 |
| year | 0.0260 | 0.0157 | 1.66 | 0.1121 |

Table 9: **Model Output:proportion of herpetofauna studies overtime.** Quasibinomial regres- sion shows no significant relationship between the proportion of studies and year.

|  | Estimate | Std. Error | t value | Pr(*>\|*t*\|*) |
| --- | --- | --- | --- | --- |
| (Intercept) | 51.4989 | 38.6132 | 1.33 | 0.1966 |
| year | -0.0267 | 0.0192 | -1.39 | 0.1780 |

Table 10: **Model Output:proportion of microbe studies overtime.** Quasibinomial regression shows no significant relationship between the proportion of studies and year.

|  | Estimate | Std. Error | z value | Pr(*>\|*z*\|*) |
| --- | --- | --- | --- | --- |
| (Intercept) | 27.4112 | 25.0745 | 1.09 | 0.2743 |
| year | -0.0146 | 0.0124 | -1.17 | 0.2405 |

Table 11: **Model Output:proportion of terrestrial mammal studies overtime.** Binomial re- gression shows no significant relationship between the proportion of studies and year.

|  | Estimate | Std. Error | z value | Pr(*>\|*z*\|*) |
| --- | --- | --- | --- | --- |
| (Intercept) | 21.7935 | 24.5123 | 0.89 | 0.3740 |
| year | -0.0118 | 0.0122 | -0.97 | 0.3328 |

Table 12: **Model Output:proportion of plant studies overtime.** Binomial regression shows no significant relationship between the proportion of studies and year.

5 **Methods model outputs**

|  | Estimate | Std. Error | z value | Pr(*>\|*z*\|*) |
| --- | --- | --- | --- | --- |
| (Intercept) | -25.8260 | 44.8392 | -0.58 | 0.5646 |
| year | 0.0112 | 0.0222 | 0.50 | 0.6144 |

Table 13: **Model Output:proportion of acoustics studies overtime.** Binomial regression shows no significant relationship between the proportion of studies and year.

| Estimate | Std. Error | z value | Pr(*>\|*z*\|*) |
| --- | --- | --- | --- |
| (Intercept) -159.2959 | 43.3793 | -3.67 | 0.0002 |
| year 0.0776 | 0.0215 | 3.61 | 0.0003 |

Table 14: **Model Output:proportion of camera trap studies overtime.** Binomial regression shows a significant relationship between the proportion of studies and year.

|  | Estimate | Std. Error | z value | Pr(*>\|*z*\|*) |
| --- | --- | --- | --- | --- |
| (Intercept) | -1.8622 | 0.1174 | -15.87 | 0.0000 |
| poly(year, 2)1 | 2.1782 | 0.6601 | 3.30 | 0.0010 |
| poly(year, 2)2 | -1.2682 | 0.4163 | -3.05 | 0.0023 |

Table 15: **Model Output:proportion of eDNA studies overtime.** Binomial regression with a quadratic term for year shows a significant curvelinear relationship between the proportion of studies and year.

|  | Estimate | Std. Error | z value | Pr(*>\|*z*\|*) |
| --- | --- | --- | --- | --- |
| (Intercept) | 66.1412 | 85.1570 | 0.78 | 0.4373 |
| year | -0.0352 | 0.0423 | -0.83 | 0.4048 |

Table 16: **Model Output:proportion of light trap studies overtime.** Binomial regression shows no significant relationship between the proportion of studies and year.

|  | Estimate | Std. Error | z value | Pr(*>\|*z*\|*) |
| --- | --- | --- | --- | --- |
| (Intercept) | 2.2359 | 39.4519 | 0.06 | 0.9548 |
| year | -0.0026 | 0.0196 | -0.13 | 0.8937 |

Table 17: **Model Output:proportion of mammal trap studies overtime.** Binomial regression shows no significant relationship between the proportion of studies and year.

|  | Estimate | Std. Error | z value | Pr(*>\|*z*\|*) |
| --- | --- | --- | --- | --- |
| (Intercept) | -2.4003 | 0.1247 | -19.25 | 0.0000 |
| poly(year, 2)1 | -0.2445 | 0.6999 | -0.35 | 0.7268 |
| poly(year, 2)2 | -1.7698 | 0.5148 | -3.44 | 0.0006 |

Table 18: **Model Output:proportion of marine transect studies overtime.** Binomial regression with a quadratic term for year shows a significant curvelinear relationship between the proportion of studies and year.

|  | Estimate | Std. Error | z value | Pr(*>\|*z*\|*) |
| --- | --- | --- | --- | --- |
| (Intercept) | 123.2629 | 33.1669 | 3.72 | 0.0002 |
| year | -0.0626 | 0.0165 | -3.80 | 0.0001 |

Table 19: **Model Output:proportion of pitfall studies overtime.** Binomial regression with a quadratic term for year shows a significant curvelinear relationship between the proportion of studies and year.

|  | Estimate | Std. Error | z value | Pr(*>\|*z*\|*) |
| --- | --- | --- | --- | --- |
| (Intercept) | -1.6752 | 0.0874 | -19.16 | 0.0000 |
| poly(year, 2)1 | -1.1795 | 0.4298 | -2.74 | 0.0061 |
| poly(year, 2)2 | 1.6879 | 0.3341 | 5.05 | 0.0000 |

Table 20: **Model Output:proportion of point count studies overtime.** Binomial regression with a quadratic term for year shows a significant curvelinear relationship between the proportion of studies and year.

|  | Estimate | Std. Error | z value | Pr(*>\|*z*\|*) |
| --- | --- | --- | --- | --- |
| (Intercept) | 58.6485 | 25.0948 | 2.34 | 0.0194 |
| year | -0.0301 | 0.0124 | -2.42 | 0.0155 |

Table 21: **Model Output:proportion of quadrat studies overtime.** Binomial regression shows a significant relationship between the proportion of studies and year.

|  | Estimate | Std. Error | z value | Pr(*>\|*z*\|*) |
| --- | --- | --- | --- | --- |
| (Intercept) | 1.7369 | 109.5742 | 0.02 | 0.9874 |
| year | -0.0034 | 0.0543 | -0.06 | 0.9498 |

Table 22: **Model Output:proportion of remote sensing studies overtime.** Binomial regression shows no significant relationship between the proportion of studies and year.

| Estimate | Std. Error | z value | Pr(*>\|*z*\|*) |
| --- | --- | --- | --- |
| (Intercept) -157.2100 | 51.7075 | -3.04 | 0.0024 |
| year 0.0764 | 0.0256 | 2.98 | 0.0029 |

Table 23: **Model Output:proportion of SCUBA studies overtime.** Binomial regression with a quadratic term for year shows a significant curvelinear relationship between the proportion of studies and year.

|  | Estimate | Std. Error | z value | Pr(*>\|*z*\|*) |
| --- | --- | --- | --- | --- |
| (Intercept) | 11.8543 | 28.8725 | 0.41 | 0.6814 |
| year | -0.0070 | 0.0143 | -0.49 | 0.6230 |

Table 24: **Model Output:proportion of terrestrial transect studies overtime.** Binomial re- gression shows no significant relationship between the proportion of studies and year.

|  | Estimate | Std. Error | z value | Pr(*>\|*z*\|*) |
| --- | --- | --- | --- | --- |
| (Intercept) | 26.0715 | 43.6013 | 0.60 | 0.5499 |
| year | -0.0146 | 0.0216 | -0.67 | 0.5003 |

Table 25: **Model Output:proportion of autonomous vehicle studies overtime.** Binomial regression shows no significant relationship between the proportion of studies and year.

|  | Estimate | Std. Error | z value | Pr(*>\|*z*\|*) |
| --- | --- | --- | --- | --- |
| (Intercept) | -31.4168 | 26.7150 | -1.18 | 0.2396 |
| year | 0.0146 | 0.0132 | 1.10 | 0.2712 |

Table 26: **Model Output:proportion of light trap studies overtime.** Binomial regression shows no significant relationship between the proportion of studies and year.

|  | Estimate | Std. Error | z value | Pr(*>\|*z*\|*) |
| --- | --- | --- | --- | --- |
| (Intercept) | -1.1264 | 0.0911 | -12.37 | 0.0000 |
| poly(year, 2)1 | 2.3556 | 0.5101 | 4.62 | 0.0000 |
| poly(year, 2)2 | -1.1974 | 0.3284 | -3.65 | 0.0003 |

Table 27: **Model Output:Change in high throughput methods over time.** Binomial regression with a quadratic term for year.

|  | Estimate | Std. Error | z value | Pr(*>\|*z*\|*) |
| --- | --- | --- | --- | --- |
| (Intercept) | -1.1654 | 0.0445 | -26.16 | 0.0000 |
| throughputLow.Total | 0.7918 | 0.0561 | 14.13 | 0.0000 |

Table 28: **Model Output:Difference between low and high throughput methods.** Binomial regression.
